# Trans-presentation of IL-15 by IL15Rα attenuates tumor immune surveillance and is dispensable for IL-15-dependent tumor growth control

**DOI:** 10.64898/2026.06.30.732683

**Authors:** Fjolla Rexhepi, Sara Ali Akbari, Mohammad Moradzad, Saeed Khodayari, Akhil Shukla, Elodie Demontier, Jean-François Lucier, Anny Armas Cayarga, Hugues Allard-Chamard, Subburaj Ilangumaran, Sheela Ramanathan

## Abstract

**Background:** IL-15 is a promising cytokine for cancer immunotherapy. IL-15 promotes differentiation and homeostasis of innate and adaptive immune cells, and their cytolytic effector functions. IL-15 receptor is composed of IL-15Rα, IL-15Rβ and the common γ_c_ chains. IL-15 is also trans-presented as IL-15Rα:IL-15 complex to IL-15Rβ:γ_c_ on neighboring cells. IL-15Rα is dispensable for early immune response to infections and in autoimmune diabetes. The role of IL-15Rα in antitumor immune responses remains unclear.

**Methods:** In WT, *Il15^−/−^* and *Il15ra^−/−^* mice, we studied the growth of syngeneic tumor cell lines and tumor immune surveillance against endogenous fibrosarcoma induced by methylcholanthrene (MCA). Immune gene signature and total proteome analysis were performed on MCA-induced tumors.

**Results:** Lack of IL-15 or IL-15Rα did not enhance the growth of implanted tumor cell lines, despite reduced immune cell infiltration. MCA-induced tumor incidence was reduced in mice lacking IL-15Rα but not IL-15, although both are required for efficient tumor immunoediting. *Il15^−/−^* and *Il15ra^−/−^* tumors showed reduced *Ifng* expression but displayed differential modulation of *Ifng-*responsive genes. Proteome profiles of *Il15^−/−^* tumors, but not tumor-derived cell lines, showed significant reduction of antigen presentation pathways. B16-F10 melanoma cells expressing NLRC5, the IFNγ-induced transcriptional activator of tumor antigen presentation, still required IL-15 but not IL-15Rα for efficient tumor control.

**Conclusions:** Our findings show that IL-15 plays a negligible role in immunosurveillance against spontaneous tumor development, whereas IL-15Rα restrains immunosurveillance. Neither IL-15 nor IL-15Rα have a significant impact on implanted tumor models, although IL-15 facilitates efficient control of highly immunogenic tumors for which IL-15Rα is dispensable.

## 1. Introduction

The IL-15 receptor (IL-15R) is composed of ligand-binding IL-15Rα and signal-transducing IL-15Rβ and γ_c_ chains, the latter two being shared with IL-2. In producer cells, IL-15 binds to IL-15Rα during biosynthesis and this complex is ‘*trans*-presented’ to responder cells expressing the IL-15Rβγ_c._ This ‘*trans’* signaling is needed for the activation of CD8^+^ T cells, NK cells and other IL-15-dependent immune cell subsets. *Il15^−/−^* mice are susceptible to viral, fungal and bacterial infections [1–7]. IL-15 has a pathogenic role in auto-immune diseases [8–13]. While many studies using IL-15 deficient mice showed the importance of IL-15 in protective immune responses during infections, very few studies directly explored the role of IL-15 trans-presentation by IL-15Rα in this protection. Unlike IL-15 deficient mice, IL-15Rα deficient mice were resistant to infections [5, 14]. Similarly, while IL-15 deficiency protected from certain autoimmune diseases, the absence of IL-15Rα did not confer protection [5, 15, 16].

Given the ability of IL-15 to activate the cytotoxicity of CD8^+^ T, NK, ILC1s and γδ T cells, IL-15 is a promising cancer immunotherapeutic agent that can enhance anti-tumor immunity. Transgenic expression of IL-15 contributes to the control of B16 tumors through the activation of NK and CD8^+^ T cells [17, 18]. MHC-I expression on tumor cells promote IL-15-dependent, CD8^+^ T cell mediated tumor control [17, 19]. Overexpressed IL-15 controlled tumor progression in genetic models of oncogene-induced tumors [20, 21]. In line with the role of NK cells in controlling tumor metastasis and hematopoietic cancers but not solid tumors [22], IL-15 deficient mice exhibited increased metastasis while transgenic expression of IL-15 diminished metastasis [18, 20]. Despite the promising anti-tumor efficacy of IL-15 predicted from its biological roles of activating cytotoxicity in NK and CD8^+^ T cells, incorporation of IL-15 in tumor immunotherapy regimens has been plagued by toxicity issues and insufficient responses in solid tumors [23]. Based on promising results obtained with a combination of BCG and IL-15 superagonist, Nogapendekin alfa inbakicept (NAI), also known as N-803 [24], N-803 has been approved by FDA for bladder cancer in 2024.

While the role of IL-15 in promoting anti-tumor responses by activating CD8+ T cells and NK cells are well established, it is not clear whether IL-15 and trans-presented IL-15 play different roles in tumor immunosurveillance. In this study we analyzed the role of IL-15 and trans-presented IL-15 in Methylcholanthrene (MCA)-induced carcinogenesis. Our observations show that incidence of tumors is comparable between WT and *Il15^−/−^* mice, while *Il15ra^−/−^* mice had reduced incidence of MCA-induced tumors. Furthermore, we show that increasing the immunogenicity of B16-F10 cells by expressing NLRC5 [25] that results in better clearance of the tumors, required IL-15 but not the trans-presented IL-15.

## 2. Materials and Methods

### 2.1. Mouse strains

Mice strains used in this study were bred in our animal facility to avoid major differences in microbiome and environmental influences. Wild type C57Bl/6 (WT) mice, *Il15^−/−^*and *Il15ra^−/−^* in the C57Bl/6 background have been described previously [5, 14, 26–28]. All experimental protocols on mice were approved by Université de Sherbrooke Ethics Committee for Animal Care and Use in accordance with guidelines established by the Canadian Council on Animal Care (protocols #2021-3144 and 2026-5225).

### 2.2. Cell lines and tumor growth

EL4, a lymphoma cell line, EG7-OVA (EG7) a lymphoma cell line derived from EL4 and expressing Ovalbumin (OVA) and B16-F10 murine melanoma cells were obtained from American Type Culture Collection (Virginia, USA). EL4 and EG7 cell lines were cultured in Roswell Park Memorial Institute (RPMI, Sigma-Aldrich) medium containing 10 % heat-inactivated fetal calf serum, 1% penicillin/streptomycin cocktail, 1% HEPES, 1% Sodium Pyruvate. For EG7-OVA, 0.4 mg/ml G418 was added to media. B16-F10 melanoma cell line was cultured in Dulbecco’s modified eagle medium (DMEM, Sigma-Aldrich) supplemented with 5% heat-inactivated fetal calf serum and 1% penicillin/streptomycin cocktail. B16-F10 expressing NLRC5 protein (reference: Galaxia) were maintained with the addition of blasticidin (8 μg/ml). All cells were grown in a humidified atmosphere of 5 % CO_2_ at 37°C.

For tumor growth 2.5x10^5^ cells in 50 μl of phosphate buffered saline (PBS) PBS were injected subcutaneously in the right hind flank of WT, *Il15^−/−^* and *Il15ra^−/−^* mice. Tumor dimensions (length and width) were recorded every three days using a digital Vernier caliper. Tumor volume was calculated using the ellipsoid formula: V = ½ x Length x Width^2^. At the endpoint (2 cm in diameter in any one direction), mice were euthanized, tumor samples were collected and analyzed for histopathological features.

### 2.3. MCA-induced fibrosarcoma for tumor immunosurveillance

Groups of WT, *Il15^−/−^* and *Il15ra^−/−^* mice were injected subcutaneously in the right hind flank with 100, 200 or 400 µg of 3-Methylcholanthrene (MCA, Sigma-Aldrich, Cat # 212942) in 50 µl of corn oil as previously described [29, 30]. The growth of MCA-induced fibrosarcoma was monitored daily over the course of 80–200 days and measured by a digital Vernier caliper along the perpendicular axes of the tumors. Tumors > 2 mm in diameter and demonstrating progressive growth were recorded as positive. Tumor samples were collected for histopathological, immunofluorescence and proteomics as previously described [30]. Tumor samples were also collected to generate MCA-induced cell lines.

### 2.4. MCA-induced fibrosarcoma cell lines and cancer immunoediting

MCA-induced fibrosarcoma were collected from WT, *Il15^−/−^* and *Il15ra^−/−^* mice. Tumor tissues were washed, minced and mechanically dissociated using gentleMACS C tubes (Miltenyi Biotec) and gentleMACS Dissociator and then incubated with Collagenase IV (1mg/ml) and DNase I (40 ug/ml) for 1 hour at 37°C for chemical digestion. Digested tissue was passed through a 40 µm filter and the collected cells were washed with PBS. Cells were cultured in RPMI medium containing 10 % heat-inactivated fetal calf serum, 1% penicillin/streptomycin cocktail, 1% HEPES, 1% Sodium Pyruvate for three months in a humidified atmosphere of 5 % CO_2_ at 37°C. Cell proliferation was assessed using Cell Counting Kit-8 (Dojindo Laboratories, Japan). For immunoediting assays *in vivo*, 2 x10^5^ MCA tumor-derived cells from WT, *Il15^−/−^* and *Il15ra^−/−^* mice were implanted subcutaneously in naive WT, *Il15^−/−^* and *Il15ra^−/−^* mice. Tumor growth was monitored from the tumor onset until the ethical limits were reached. Tumor samples were collected and analyzed as described above.

### 2.5. RNA isolation and gene expression analysis

RNA from frozen WT, *Il15^−/−^* and *Il15ra^−/−^* tumors and MCA-derived cell lines was extracted using RNeasy Mini kit (Qiagen, Toronto, Ontario, Canada. cDNA was synthetized from 500ng of purified RNA using QuantiTect Reverse Transcription Kit (Qiagen, Toronto, Ontario, Canada). Quantitative RT-PCR amplification reactions were carried out in QuantStudio 3 Real-Time PCR System (Thermo Fisher Scientific, Canada) using SYBR Green Supermix (Bio-Rad, Mississauga, Ontario, Canada). The expression of indicated genes was measured using primers listed in Supplementary Table S1. Gene expression levels between samples were normalized based on the Cycle threshold (Ct) values compared to housekeeping gene 36B4 (*Rplp0*).

### 2.6. Histopathological and Immunofluorescence analysis

Formalin-fixed paraffin-embedded tissue sections of 4 μm thickness were deparaffinized, rehydrated, and stained with hematoxylin and eosin (H&E). Digital images of the stained sections were acquired using a Nanozoomer Slide Scanner (Hamamatsu Photonics, Japan) and analyzed by the Nanozoomer Digital Pathology software NDPview2.7.52 (Hamamatsu Photonics). For immunofluorescence, deparaffinized and rehydrated tumor sections were incubated in Buffer A (Electron Microscopy Sciences, #62707-10) for antigen unmasking using Retriever 2100 (Electron Microscopy Sciences, #R2100-US). Tumor sections were then blocked using 5% BSA in Tris-bu/ered saline (TBS) containing 20% Tween-20 (TBS-T) followed by overnight incubation with primary antibody at 4°C. After washing, tissue sections were incubated with appropriate secondary antibody for 2h, washed and nuclei were stained using Hoechst 33342 DNA staining dye (Thermo Fisher Scientific, Canada). After washing, tissue sections were mounted with a coverslip with Dako Fluorescence Mounting Media (cat#S3023) and images were taken using fluorescent microscope Zeiss Axioscope 2.

### 2.7. Mass spectrometry

The MCA tumor samples and tumor-derived cell lines were processed for proteomic analysis as previously described [30]. Briefly, 25 to 50 mg of frozen tumors were lysed with a Lysis buffer (5% sodium dodecyl sulfate (SDS), 50 mM triethylammonium bicarbonate (TEAB), pH 8.5), incubated at 95°C for 3 minutes and homogenized using Qiagen TissueLyser (Qiagen). The DNA was sheared by sonication before protein quantification (Pierce BCA Protein Assay Kit, ThermoFisher, ref#23227). 100ug of proteins were reduced in 100 μl of Lysis buffer by adding Tris(2-carboxyethyl)phosphine (TCEP) to 5mM final concentration, followed by a 55°C, 15 min incubation. Proteins were alkylated by adding chloroacetamide (Sigma-Aldrich, Saint-Louis) to a final concentration of 20 mM and incubated for 10 minutes at room temperature. Complete denaturation of proteins was done by adding phosphoric acid to a final concentration of 2.5%. A binding/wash buffer was then added to protein samples to a final concentration of 100mM TEAB in 90% methanol before being transferred to a S-trap column. The column was then washed three times. Proteins were digested by adding 1 μg of Pierce mass spectrometry (MS)-grade trypsin (Thermo Fisher Scientific, Waltham) and incubated overnight at 30°C. Peptides were then eluted using 50 mM TEAB, 0.2% formic acid and 50% acetonitrile before being concentrated by centrifugal evaporation and then resuspended in 1% formic acid. Peptides were quantified using a NanoDrop spectrophotometer (Thermo Fisher Scientific, Waltham, MA).

For mass data acquisition in data independent acquisition (DIA) mode, 250 ng of peptides from each sample were injected into an HPLC (nanoElute, Bruker Daltonics) and loaded onto a trap column with a constant flow of 4 μL/min (AcclaimPepMap100 C18 column,0.3 mm id × 5 mm, Dionex Corporation), and then eluted onto an analytical C18 Column (1.9 µm beads size, 75 μm × 25 cm, PepSep) heated at 50°C. Peptides were eluted for 2 h over gradient of acetonitrile (5%–37%) in 0.1% FA at 400 nL/min while being injected into a TimsTOF Pro ion mobility mass spectrometer equipped with a Captive Spray nano electrospray source (Bruker Daltonics). Data was acquired using diaPASEF mode (Meier et al., 2020). Briefly, for each single Trapped Ion Mobility Spectrometry (TIMS; 100 ms) in diaPASEF mode, 1 mobility window consisting of 27 mass steps (m/z between 114 and 1,414 with amass width of 50 Da) per cycle (1.27 sec duty cycle) was used. These steps cover the diagonal scan line for +2 and +3 charged peptides in them/z-ion mobility plane.

### 2.8. Proteomics data analysis

Peptide mass spectra were analyzed using the DIA-NN, an open-source software [31] suite for DIA / SWATH data processing (https://github.com/vdemichev/DiaNN, version 1.8.1), installed in an Apptainer container (https://apptainer.org/, version 1.3.5) using docker image provided on the docker hub [32]. Analysis was performed using default parameters, except for the following options: two missed cleavages were allowed; trypsin digestion was performed for K/R; and protein N-term methionine excision was used as a variable modification for the in-silico digest as described [30].

The Mus musculus reference proteome UP000000589 was downloaded from the UniProt website (https://www.uniprot.org/proteomes/UP000000589). The reference proteome contained 63367 proteins. For the FASTA search, DIA-NN was instructed to perform an in silico digest of the sequence database. A mass tolerance accuracy of MS1 and MS2 of 20 ppm was used for the precursor and fragment ions, respectively. Minimum and maximum values were set for peptide length (7-30 amino acids), precursor charge (1-5), precursor m/z (100-1700) and fragmentation m/z (100-1500) for in silico library generation or library-free search. For re-analysis, match between runs (MBR) was enabled and smart profiling was chosen when creating a spectral library from DIA data. Carboxyamidomethylation (unimod4) and oxidation (M) (unimod35) were set as fixed modifications, and N-terminal protein acetylation was set as a variable modification. DIANN protein group matrix was filtered using a custom Perl script to extract protein groups with a single protein.

The proteomic data was analyzed using in-house customized R (4.6.0) scripts for data import, data quality check, analysis of differential proteins expression using the Limma package, generation of volcano plots, Gene Set Enrichment Analysis (GSEA) using KEGG and Hallmark gene sets, and pathway enrichment heatmaps. The complete analysis workflow is available on GitHub (https://github.com/MohammadMoradzad/tumor-proteomics-analysis-pipeline).

### 2.9. Flow cytometry

Tumor infiltrating lymphocytes (TILs) were isolated using Tumor Dissociation Kit, mouse (#130-096- 730) from Miltenyi Biotec, Germany. Single cell suspensions of TILs, draining lymph nodes (DLN) and non-draining lymph nodes (NDLN) were washed with PBS containing 1% FCS and aliquoted up to 1 x 10^6^ cells/100μl into FACS tubes for antibody staining for flow cytometry. The cells were labeled with fluorochrome-conjugated antibodies for flow cytometry as described [33]. Cells were analyzed on a Cytoflex 30 flow cytometer (Beckman Coulter, Brea, CA, USA) or BD FACSDiscover^TM^ S8 cell sorter (BD Biosciences, CA, USA) and data analyzed using the FlowJo software versions 10 or 11 (BD Biosciences). The list of antibodies used is given in Supplementary Table S2 for Cytoflex 30 and Supplementary Table S3 for BD FACSDiscover^TM^ S8.

### 2.10. Data Analysis

Graphpad Prism software (Version 10) was used for data analysis and to generate illustrations. The data are shown as mean ± standard deviation (SD). One- or Two-way ANOVA with Tukey’s or Bonferroni’s post-hoc test was used as indicated in figure legends to compare datasets and to calculate statistical significance.

## 3. Results

### 3.1. Lack of IL-15 or IL-15Rα does not enhance the growth of implanted syngeneic tumors

Innate immune responses that contribute to the control of viral and bacterial infections and promote autoimmunity require IL-15 signaling without the need for IL-15 trans-presentation by IL-15Rα [5, 8, 14, 16]. Even though NK cells are implicated in IL-15-mediated control of an oncogene-driven breast cancer model [20], their role in solid tumors is not clear [34]. To investigate how endogenous levels of IL-15 and its trans-presentation via IL-15Rα impact tumor growth, we implanted in wild type (WT), *Il15^−/−^* and *Il15ra^−/−^* mice syngeneic tumor cell lines that vary in MHC-I expression and in the ability to activate antitumor immune cell subsets. EL4 is a T-cell lymphoma that expresses high levels of MHC-I and EG7 is an EL4-derived line expressing chicken ovalbumin as a surrogate tumor antigen to render it highly immunogenic [35]. All control (wild type, WT), *Il15^−/−^* and *Il15ra^−/−^*mice implanted with EL4 developed tumors between 7- and 19-days post-implantation with no significant differences in tumor size or volume (Supplementary Figure. S1A). EG7 cells elicit a strong CD8^+^ T cell response resulting in efficient tumor control, requiring larger inoculum (>3 x 10^6^ cells) for tumor formation following subcutaneous implantation [36]. To determine if the loss of IL-15 or IL-15Rα would allow tumor growth by sub-optimal dose of EG7, 3 x 10^5^ cells were implanted in WT, *Il15^−/−^* and *Il15ra^−/−^* mice. Whereas 3/7 WT (57%), 1/4 *Il15^−/−^* (25%) and 3/6 *Il15ra^−/−^*(50%) mice developed tumors between 22 and 50 days, the rest remained tumor-free (Supplementary Fig. S1B), indicating that loss of IL-15 or IL-15Rα did not increase tumor formation by EG7 cells. B16-F10 melanoma cells express low levels MHC-I and are inefficient in activating CD8^+^ T cells but can be targeted by NK cell-mediated killing [37, 38]. As trans-presented IL-15 is required for NK cell homeostasis [39], we assessed the growth of B16-F10 melanoma in WT, *Il15^−/−^* and *Il15ra^−/−^*mice. The growth of B16-F10 melanoma in *Il15^−/−^* and *Il15ra^−/−^* mice was comparable to that in WT mice (Supplementary Fig. S1C), even though the number of NK cells within tumor infiltrating leukocytes (TILs) was significantly reduced in *Il15^−/−^* mice (Supplementary Figure. S2). NKT cells, CD8^+^ T cell subsets and MHC class-II^+^ dendritic cells were also significantly reduced in B16-F10 tumor draining lymph nodes (DLN) of *Il15^−/−^* mice (Supplementary Figure. S2), suggesting inefficient activation of these cells in the absence of IL-15. A similar pattern was observed in *Il15ra^−/−^*mice but the reduction was modest. Overall, the lack of IL-15 or IL-15Rα did not result in increased growth of B16-F10, EL4 or EG7 tumors, suggesting that these tumor cell line models may not be robust enough to investigate the impact of basal levels of IL-15 and IL-15 trans-signaling via IL-15Rα on the immune control of solid tumor development.

### 3.2. IL-15Rα restrains immune surveillance of endogenously arising tumors

As IL-15 signaling regulates innate immune responses of NK, NKT and γδ T cells, in addition to regulating some CD8^+^ T cell subsets, we assessed the requirement of IL-15 and IL-15Rα in the control endogenously arising tumors that are restrained by immune surveillance mechanisms [40]. A widely used model of tumor immune surveillance is the 3-methylcholanthrene (MCA)-induced fibrosarcoma, which demonstrated the requirement of CD8^+^ T cells and the effector molecules perforin and granzyme B for tumor immune surveillance [41–43]. Earlier studies using anti-NK1.1 and anti-NKG2D antibodies indicated a requirement for NK and NKT cells in the immune surveillance of MCA-induced tumors [44, 45]. However, using mice with targeted deletion of NK and ILC1s it was shown that NK and ILC1 cells are dispensable and even suggested a role of these cells in inhibiting immune surveillance against MCA tumors [46]. As the MCA dose required to induced MCA tumors vary with genetic background and in different mice colonies [42, 43], we used 100, 200 or 400 μg MCA in WT, *Il15^−/−^*and *Il15ra^−/−^* mice (Figure. 1A-C). Data from both sexes were pooled as male and female mice showed no significant difference in tumor incidence. Only a few mice in all three genotypes developed tumors at 100 μg MCA dose, whereas IL-15Rα deficient mice showed significantly decreased tumor incidence at 200 μg MCA dose, when compared to WT and *Il15^−/−^*mice (Fig. 1B). Cumulative tumor incidence at all MCA doses revealed that mice lacking IL-15Rα displayed resistance to MCA-induced tumor development compared to WT and *Il15^−/−^* mice (Figure. 1B,D). These results indicate that absence of IL-15 does not influence the kinetics of tumor development in the MCA model, whereas absence of IL-15 trans-presentation by IL-15Rα reduces tumor incidence.

**Figure 1.**
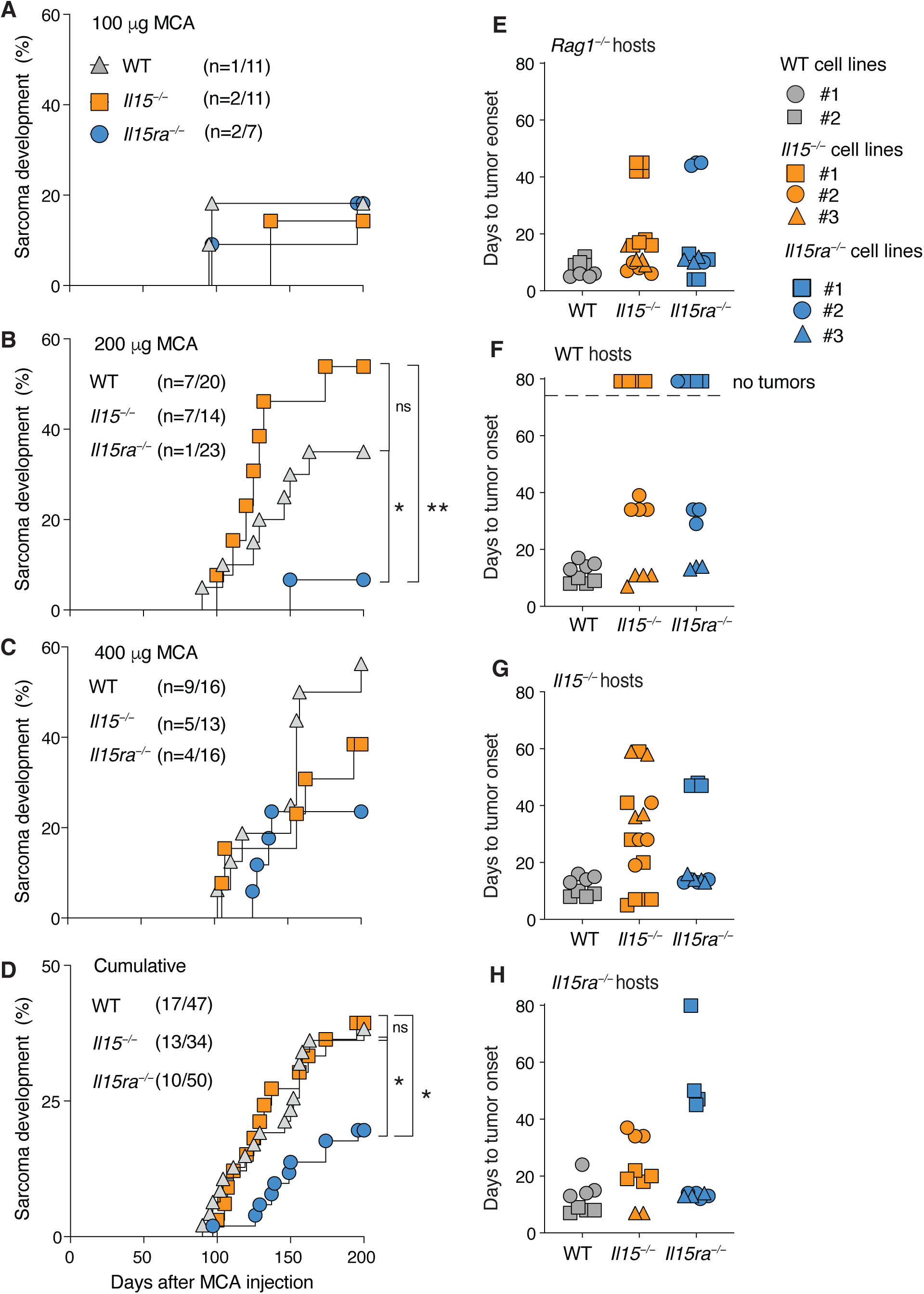
Absence of trans-presented IL-15 reduces the susceptibility to spontaneous MCA-induced fibrosarcoma. WT, *Il15^−/−^*and *Il15ra^−/−^* mice were injected the chemical carcinogen 3-ethylcholanthrene (MCA) subcutaneously. Development of fibrosarcoma was monitored visually and by using a digital vernier caliper until the endpoint (tumor diameter of 20mm in any direction, or ulceration at any diameter), when the animals were euthanized. Kaplan-Meyer curve for the number of mice that developed tumors at 100 (A), 200 (B), 400 μg (C) of MCA and (D) cumulative data. Log-rank (Mantel-Cox) test performed using GraphPad PRISM to compare survival probability. * p ≤0.05. MCA-derived tumor cell lines from WT, *Il15^−/−^*and *Il15ra^−/−^* mice were implanted in (E) *Rag1^−/−^*, (F) WT, (G) *Il15^−/−^* and (H) *Il15ra^−/−^* mice and followed for tumor onset.

### 3.3 Impact of IL-15 or IL-15Rα deficiency on tumor immunoediting

MCA-induced tumors developing in mice with impaired antitumor immune mechanisms are not efficiently immunoedited, and thus cell lines established from these tumors are effectively controlled in immune-sufficient hosts upon transplantation but grow unhindered in immunodeficient hosts [30, 47]. As MCA-induced tumor incidence was comparable between WT and *Il15^−/−^* mice but decreased in *Il15ra^−/−^* mice, we assessed whether the MCA tumors from *Il15^−/−^* and *Il15ra^−/−^* mice differed in their immunoedited status. MCA tumor cell lines established from the WT, *Il15^−/−^* and *Il15ra^−/−^* mice showed comparable growth *in vitro* and were equally responsive to IFNγ in upregulating MHC-I and PD-L1 (Supplementary Figure. S3A-C). These cell lines were then tested for *in vivo* growth in *Rag1^−/−^,* WT, *Il15^−/−^* and *Il15ra^−/−^* mice. All the tested cell lines formed tumors in *Rag1^−/−^* mice that do not have the adaptive immune system, as expected (Figure. 1E). The WT tumor cell lines rapidly formed tumors in WT, *Il15^−/−^* and *Il15ra^−/−^* recipients, confirming their immunoedited and thus immune escape status (Figure. 1F-H). Of the three cell lines derived from *Il15ra^−/−^* and *Il15^−/−^*tumors, one *Il15^−/−^* and one *Il15ra^−/−^* cell lines failed to form tumors in WT hosts indicating that they are well controlled. However, most *Il15^−/−^* and *Il15ra^−/−^*tumor cell lines formed delayed tumors in *Il15ra^−/−^* and *Il15^−/−^* hosts, suggesting that MCA tumors developing in *Il15^−/−^* and *Il15ra^−/−^* mice might not have been efficiently immune edited, possibly due to the paucity of IL-15 dependent adaptive and innate lymphocytes during tumor development and progression.

### 3.4. Immune response gene expression in MCA tumors of *Il15^−/−^* and *Il15ra^−/−^* mice

Hematoxylin and eosin staining of tumor tissues sections did not reveal any infiltrating immune cell clusters (Figure. 2A). Nonetheless, immunofluorescence staining of tumor sections indicated scattered distribution of CD45^+^ cells in all the three genotypes (Figure. 2B) but higher staining for CD31 in IL-15 deficient tumors, an endothelial cell marker for tumor angiogenesis (Fig. 2C). We analyzed the expression of select set of genes that are characteristic of antitumor immune responses. *Klrc2* (NKG2C), which is expressed on NK cells and some CD8^+^ T cell subsets [48], was reduced in both *Il15^−/−^* and *Il15ra^−/−^* tumors (Figure. 2D). On the other hand, *Klrb1c* (NK1.1) was reduced in *Il15ra^−/−^*tumors but *Il15^−/−^* and *Il15ra^−/−^* tumors, whereas *Klrk1* (NKG2D) showed an inverse expression pattern (Figure. 2D). The expression of *Klrc2* was unaffected in *Il15^−/−^* or *Il15ra^−/−^* tumors. Both *Il15^−/−^* and *Il15ra^−/−^*tumors displayed reduced C*d8* expression compared to WT tumors, whereas *Cd4* expression was unaffected (Figure. 2E). Notably, *Ifng* but not *Ifna* was greatly reduced in both *Il15^−/−^* and *Il15ra^−/−^* tumors when compared to WT tumors (Figure. 2F). The expression of *Pdl1*, which is induced by *Ifng* was reduced only in *Il15ra^−/−^*tumors, whereas the IFN-inducible *Cxcl9* and *Cxcl10* gene expression was reduced in *Il15^−/−^* tumors but not in *Il15ra^−/−^* tumors (Figure. 2G). These observations suggest that the immune responses towards MCA tumors developing in *Il15^−/−^* mice and *Il15ra^−/−^*mice share certain features but show considerable differences.

**Figure 2:**
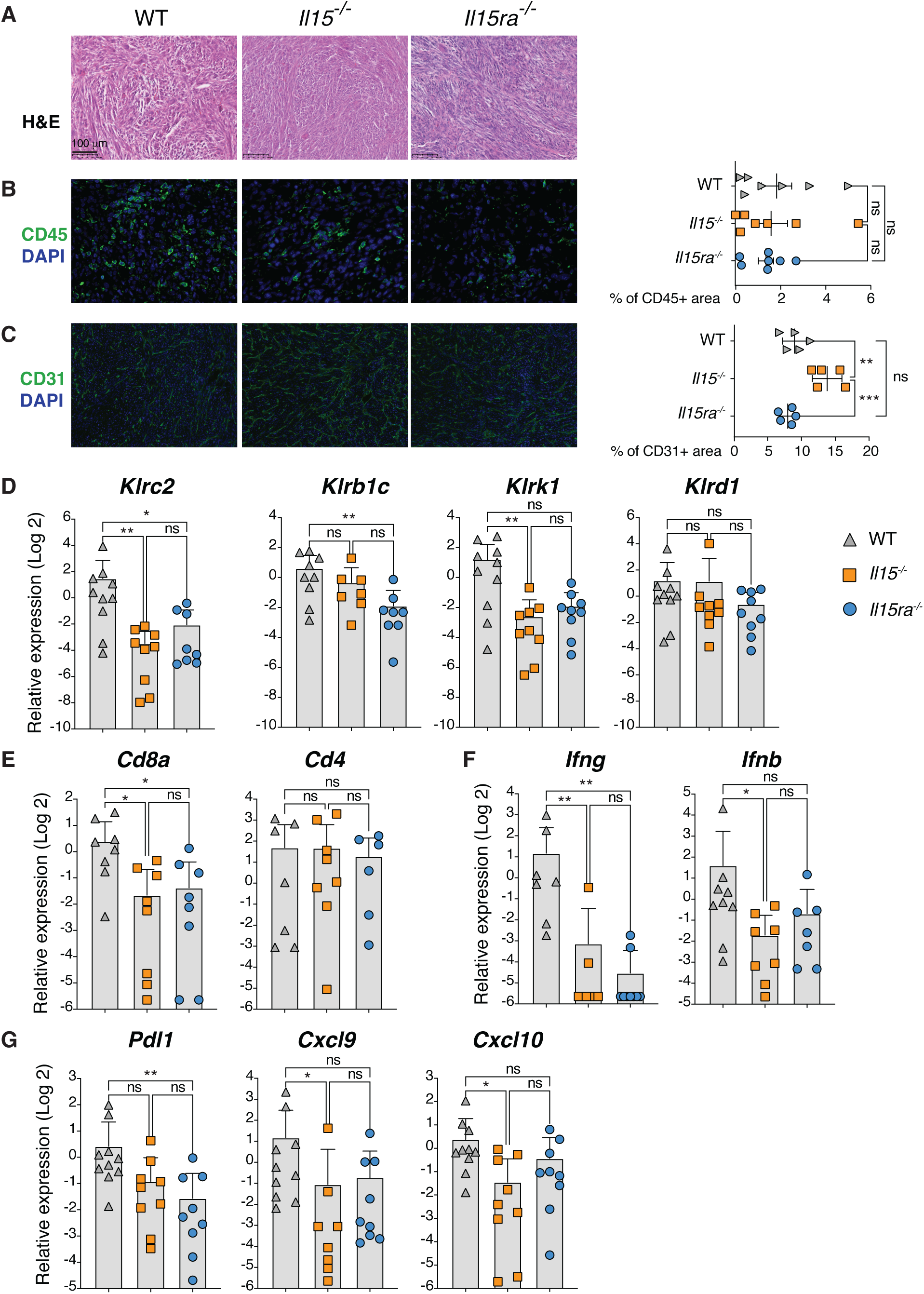
Characterization of MCA tumors. (A) Representative figures of H&E staining (20x magnification, top panel) and (B) immunofluorescence staining for CD45/DAPI (middle) and (C) CD31/DAPI (bottom) of MCA tumors are shown. The bar graphs on the right show the % staining for CD45- or CD31- positive area. Gene expression analyses in spontaneous MCA-induced fibrosarcoma was carried out on total RNA extracted from 6-8 tumors per group and analyzed by RT-qPCR for the indicated genes. (D) genes associated with NK cells, (E) *Cd4* and *Cd8*, (F) *Ifna* and *Ifng* and (G) *Ifn* stimulated genes. Statistics: Mean + SD. Ordinary one-way ANOVA with Tukey’s multiple comparison test. * *p* ≤0.05, ** *p* ≤0.01

### 3.5. Overlapping and distinct proteome profiles of *Il15^−/−^* and *Il15ra^−/−^* MCA tumors

Despite the expected heterogeneity of oncogenic drivers in MCA-induced sarcomas, somatic mutation loads in MCA-induced tumors in WT mice are lower when compared to radiation-induced tumors in WT mice [49]. Hence, we carried out proteomic analyses of MCA tumors that developed in WT, *Il15ra^−/−^* and *Il15^−/−^* mice to identify any differences in signaling pathways arising from the absence of IL-15 or IL-15Rα during tumor development. Quality control data showed 6000 to 7000 individual proteins in all samples, with comparable level of protein intensity distribution and acceptable levels of missing values in replicates (Supplementary Figure. S4A-C). Notably, principal component analysis showed clear separation of *Il15^−/−^*tumors from WT and *Il15ra^−/−^* tumors that showed partial overlap, supported by sample-to-sample correlation (Supplementary Figure. S4D-F). Volcano plots and Venn diagrams showed hundreds of differentially regulated proteins (DEPs) uniquely in tumors arising from *Il15^−/−^* and *Il15ra^−/−^* mice compared to WT tumors (Figure. 3A-D), suggesting that lack of IL-15 or IL-15Rα differentially affects protein expression across the replicates, notably, a set of proteins that are downregulated in *Il15^−/−^* tumors but upregulated in *Il15ra^−/−^* tumors and vice versa, which encompass proteins involved in general cellular homeostasis (Figure. 3E). These observations suggest that IL-15 signaling in tumors could modulate signaling pathways within tumors depending on the availability of IL-15Rα.

**Figure 3:**
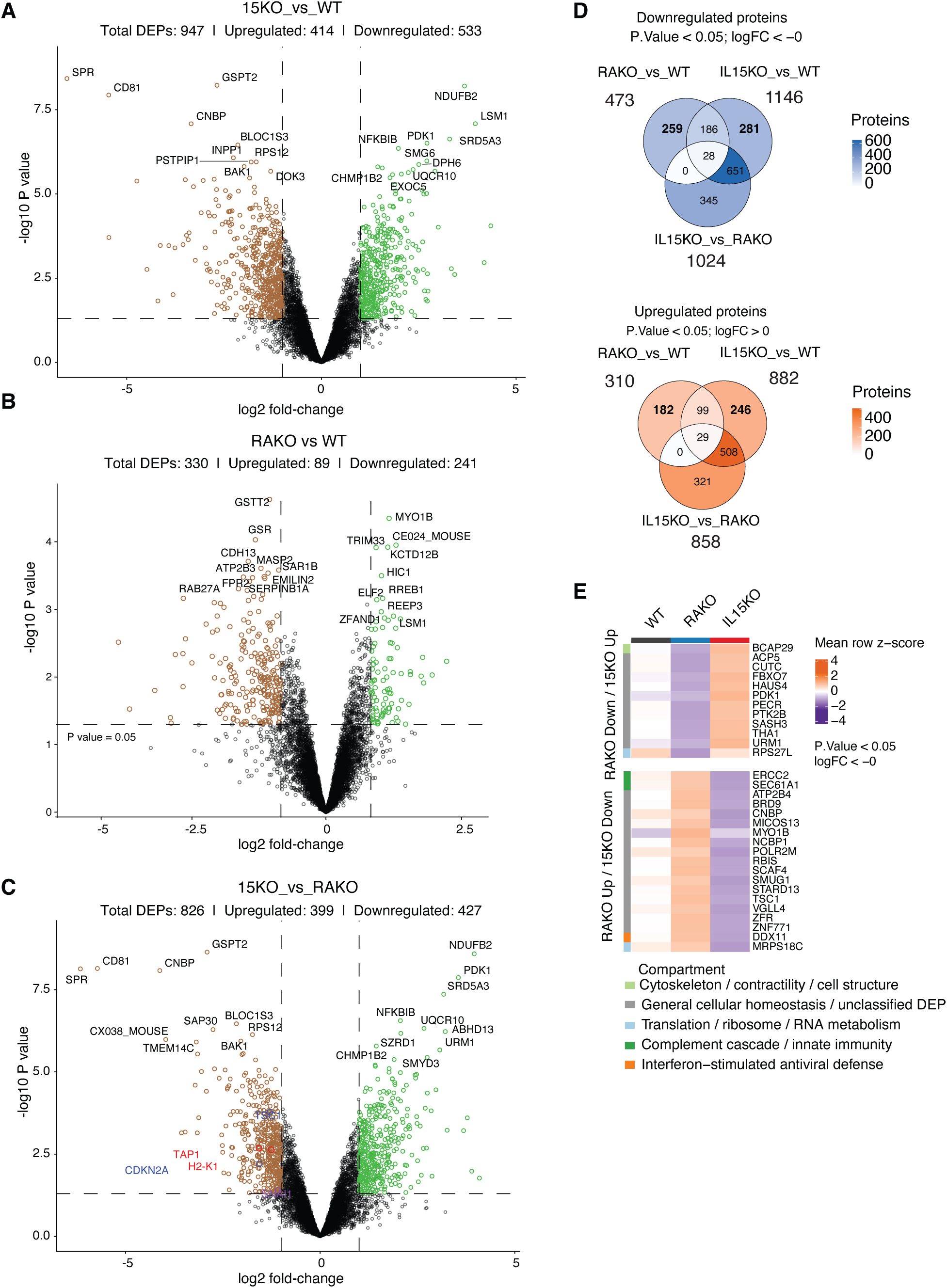
Proteomes of MCA tumors. Quadruplicate samples of proteins extracted from MCA tumors were analyzed using mass spectrometry. (A-C) Volcano plots of proteins detected in *Il15^−/−^* (15KO) tumors (A) and *Il15ra^−/−^* (RAKO) tumors (B) compared to WT tumors and 15KO tumors compared to RAKO tumors (C). Upregulated proteins are indicated in green color and downregulated proteins in brown color. (D) Venn diagrams showing the overlap between the pairwise comparisons shown in A-C within the upregulated and downregulated proteins. (E) Heat map analyses of DEPs that are differentially expressed between 15KO and RAKO tumors.

To determine how IL-15 and IL-15Rα impact signaling pathways in tumors, gene set enrichment analysis (GSEA) using Hallmark and KEGG pathways was performed on the DEPs in MCA tumors from *Il15^−/−^* and *Il15ra^−/−^*mice (Figure. 4). Among the Hallmark pathways, IFNα and IFNγ responses were significantly downregulated in both IL-15 deficient and IL-15Rα deficient tumors when compared to WT tumors (Figure. 4A-B). A heatmap analysis of proteins modulated in these two overlapping pathways is shown in Figure. 5A. IL-2-Stat5 signaling pathway, that is shared between IL-15 and IL-2 [50] was downregulated in *Il15^−/−^* tumors when compared to WT and *Il15ra^−/−^* tumors as expected, and indicating towards the possibility of IL-15Rα independent IL-15 signaling (Supplementary Figure. 5). Analysis of the KEGG pathways revealed significant downregulation of leukocyte trans-endothelial migration and antigen processing and presentation pathways in IL-15 deficient tumors when compared to WT tumors (Fig. 4D, Fig. 5B,C). Notably, IL-15Rα deficient tumors showed significant downregulation of neutrophil extracellular trap formation, and upregulation of Notch signaling pathway, polycomb repressive complex and basal transcription factors compared to WT tumors (Fig. 4D-E, 5E-F). These pathways modulated in IL-15Rα deficient tumors likely require the presence of IL-15, as they were not represented among the DEPs between *Il15^−/−^*and *Il15ra^−/−^* tumors (Fig. 4C,F). The above proteomic analyses of the MCA tumors comprise pathways associated with the tumor cells and the tumor microenvironment of the host including the immune cells. To dissociate the tumor cell-intrinsic pathways modulated by the lack of IL-15 or IL-15Rα, we performed proteome analysis of MCA tumor-derived cell lines that were grown in culture. Hallmark and KEGG pathway analyses showed that processes associated with immune responses such as interferon signaling and antigen processing and presentation were not represented among the pathways modulated in MCA tumor-derived cell lines (Supplementary Figure. 6-7). Comparison of the Hallmark and KEGG pathways modulated by IL-15 or IL-15Rα deficiency in tumor tissues and in tumor-derived cell lines revealed several shared pathways (Supplementary Figure. 8). However, downmodulation of interferon responses, antigen processing and presentation, and neutrophil extracellular traps (NET) formation were observed only in MCA tumors and not in cell lines (Figure. 6). These findings indicated that IL-15 signaling within tumor cells and the tumor microenvironment modulates host immune responses, some of which are differentially impacted by IL-15Rα-dependent regulation.

**Figure 4:**
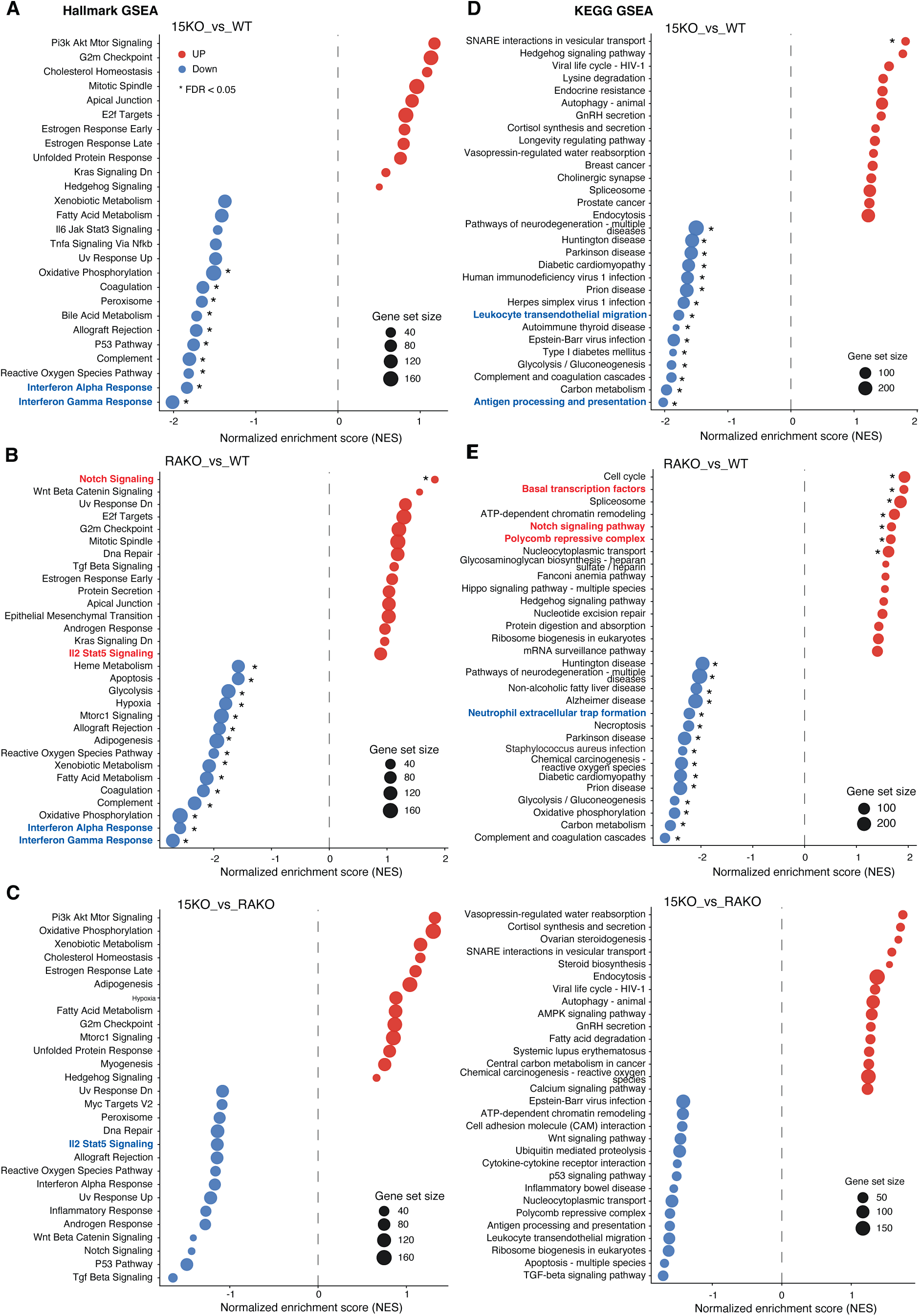
Differentially expressed pathways from proteome analyses of MCA tumors. Top 15 upregulated (in red) and downregulated (in blue) pathways that are different between 15KO tumors (A,D) and RAKO tumors (B,E) compared to WT tumors and 15KO tumors compared to RAKO tumors (C,F) by Hallmark (A-C) and KEGG (D-F) pathway analyses are shown.

**Figure 5:**
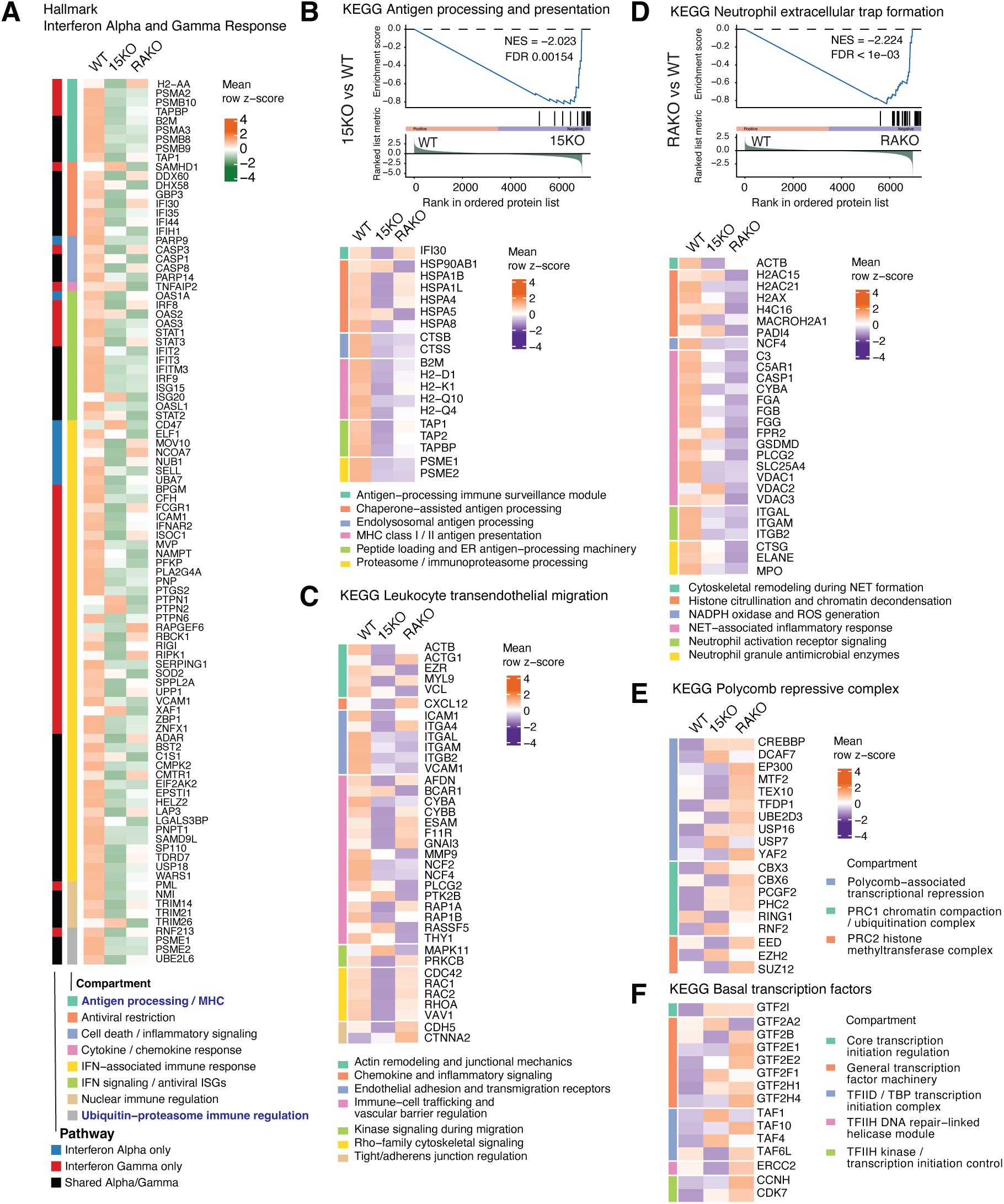
Heat map analyses of immune related differentially expressed DEPs. (A) Interferon alpha and gamma responses in Hallmark; (B) Antigen processing and presentation, (C) Leukocyte transendothelial migration-KEGG, (D) Neutrophil extracellular trap formation-KEGG, (E) Polycomb repressive complex and (F) Basal transcription factors.

**Figure 6:**
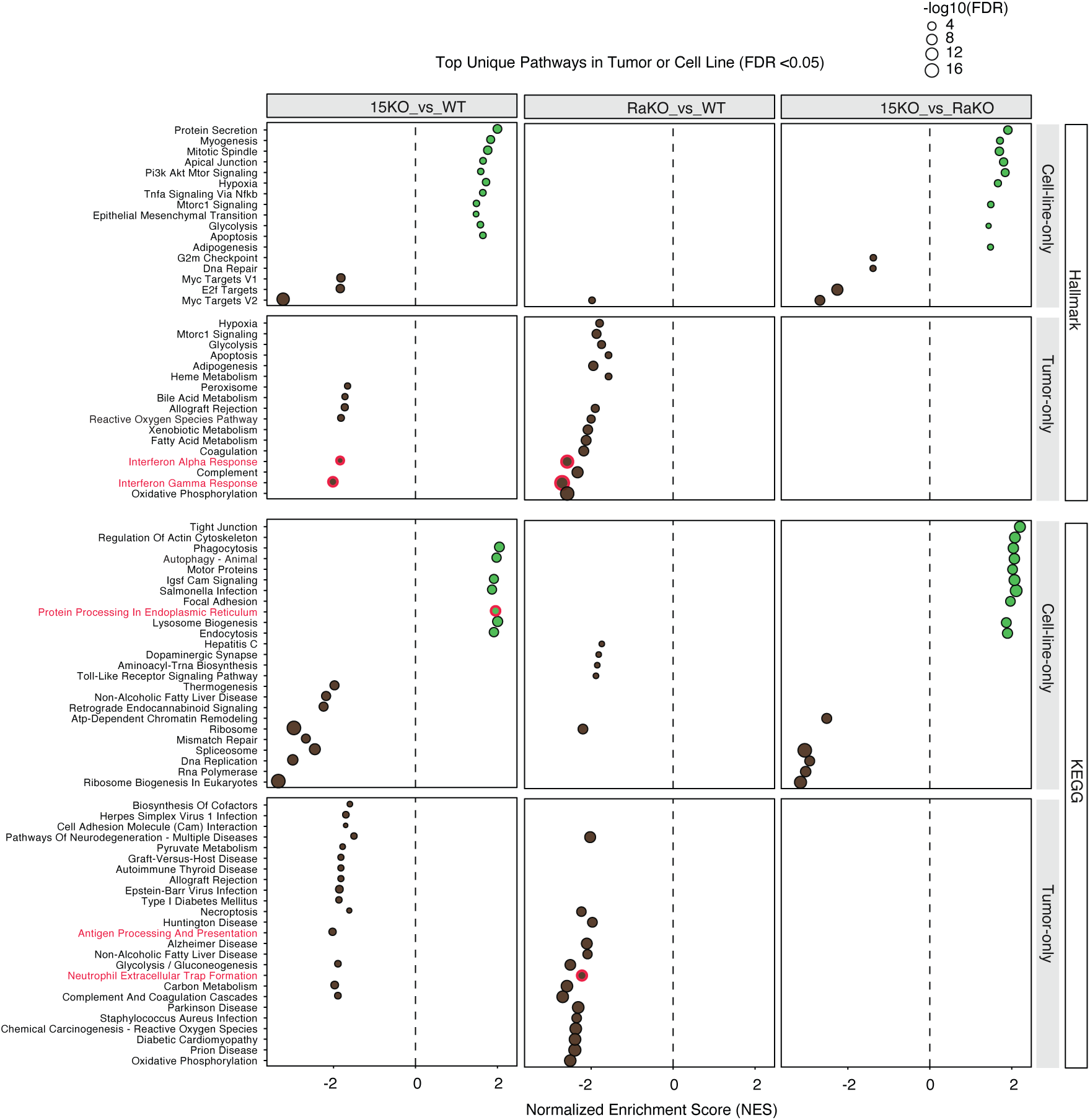
Differentially expressed pathways from proteome analyses of MCA tumor derived cell lines. Significantly upregulated (in green) and downregulated (in black) Hallmark and KEGG pathways that are unique to MCA tumor derived cell lines or tumors are shown.

**Figure 7.**
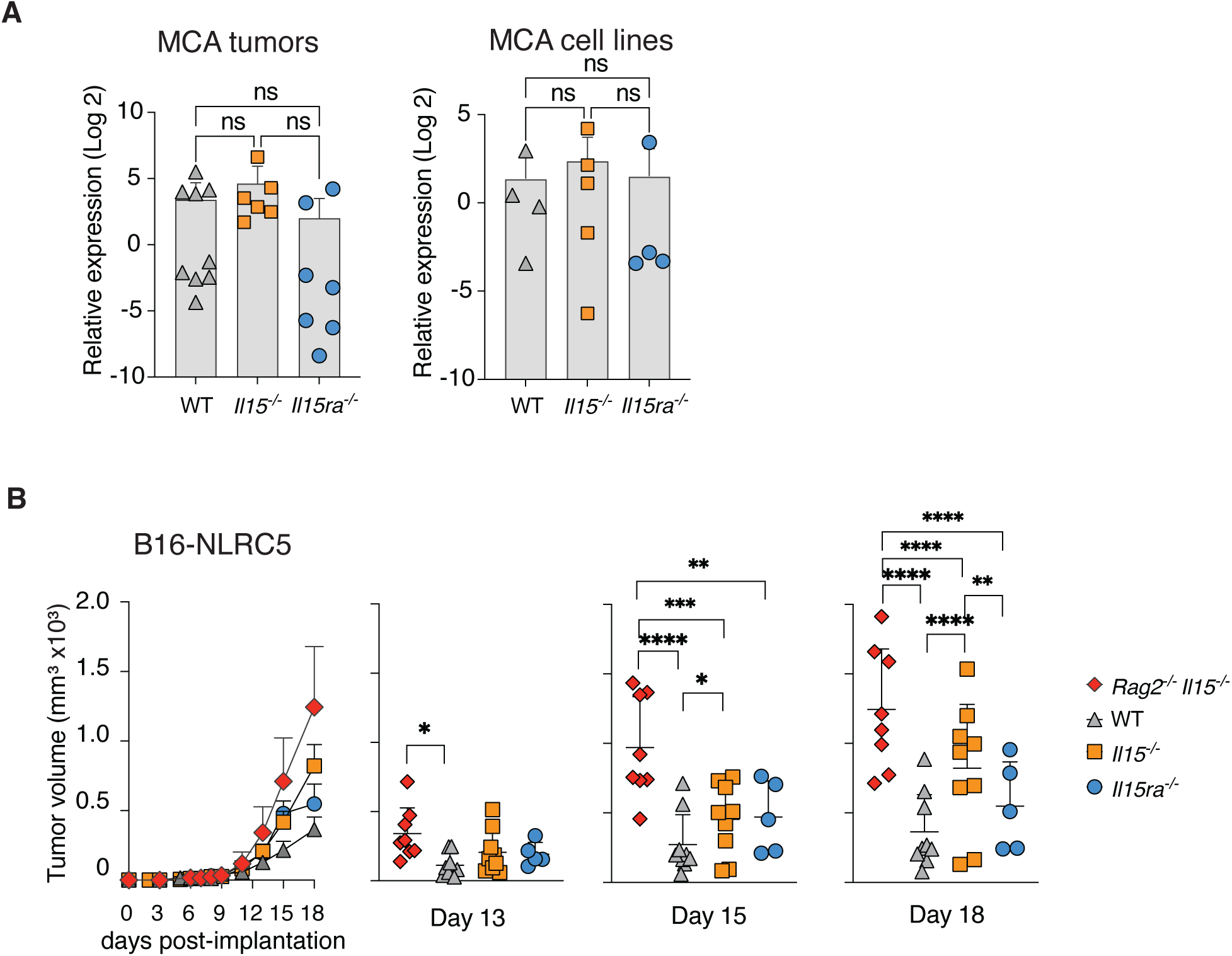
IL-15 trans-presentation by IL-15Rα is not needed for controlling B16-F10 tumors expressing NLRC5. (A) *Nlrc5* gene expression is comparable in MCA tumors and MCA-tumor derived cell lines. (B) Tumor growth in WT, *Il15^−/−^*, *Il15ra^−/−^* and *Rag1^-/-^ Il15^−/−^* mice following subcutaneous implantation of B16-NLRC5 cells. Statistics: Mean + SEM. Ordinary one-way ANOVA with Tukey’s multiple comparison test. * *p* ≤0.05, ** *p* ≤0.01, *** *p* ≤0.001, **** *p* ≤0.0001.

**Figure 8:**
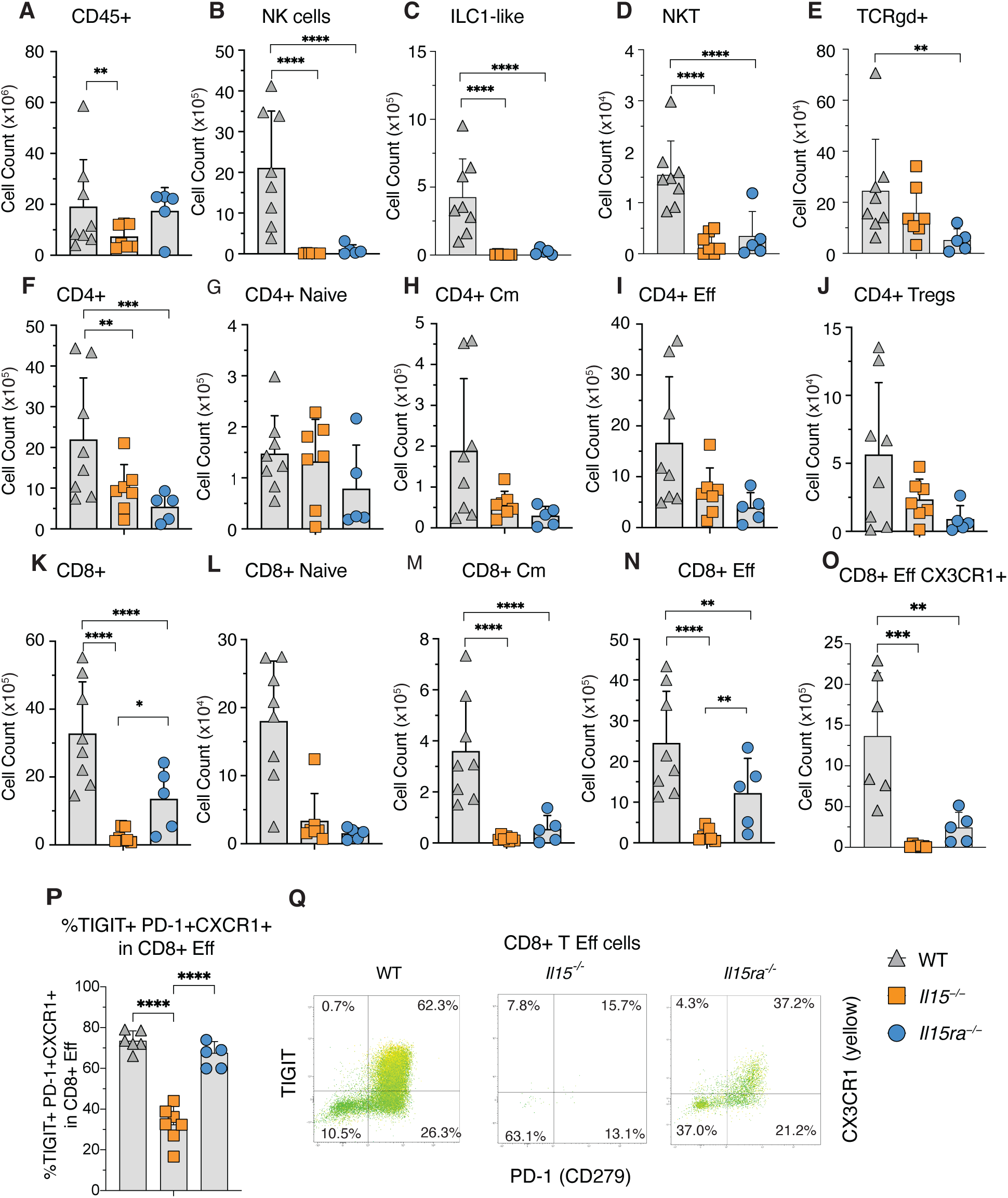
Lymphocyte subsets in TILs of B16-F10-NLRC5 tumors. Tumor infiltrating lymphocytes (TILs) were isolated at the time of tumor collection from WT, *Il15^−/−^* and *Il15ra^−/−^* mice hosts implanted with B16-NLRC5 cells. Cells were counted and stained with fluorochrome conjugated antibodies for lymphoid and myeloid cell markers and analyzed by flow cytometry using gating strategies depicted in Supplementary Fig. S9. Data from 7-9 mice per group from three independent experiments are shown. The cell numbers in TILs were equalized to the weight of the tumor mass (TILs per gram; TILS(g^-1^)). (A) Absolute number of live CD45^+^ cells, (B) Absolute number of NK cells (CD45^+^Ly6G^-^CD19^-^CD20^-^CD3^-^CD49a^-^NK1.1^+^), (C) Absolute number of ILC1-like cells (CD45^+^Ly6G^-^CD19^-^CD20^-^CD3^-^CD49a^+^NK1.1^+^), (D) Absolute number of NKT cells (CD45^+^Ly6G^-^CD19^-^CD20^-^CD3^+/-^TCRgd^-^CD4^+^NK1.1^-^), (E) Absolute number of TCRψ8 T cells (CD45^+^Ly6G^-^CD19^-^CD20^-^CD3^+/-^TCRgd^+^). (F) absolute number of CD4^+^ T cells (CD45^+^Ly6G^-^CD19^-^CD20^-^CD3^-/+^TCRgd^-^CD8^-^CD4^+^NK1.1^-^), (G) absolute number of CD4^+^ naïve T cells (CD45^+^Ly6G^-^CD19^-^CD20^-^CD3^-/+^TCRgd^-^CD8^-^CD4^+^NK1.1^-^CD25^-^CD62L^+^CD44^-^), (H) Absolute number of CD4^+^ Cm T cells (CD45^+^Ly6G^-^CD19^-^CD20^-^CD3^-/+^TCRgd^-^CD8^-^CD4^+^CD62L^+^CD44^+^), (I) absolute number of CD4^+^ Eff T cells (CD45^+^Ly6G^-^CD19^-^CD20^-^CD3^-/+^TCRgd^-^CD8^-^CD4^+^CD62L^-^CD44^+^), (J) absolute number of CD4^+^ Treg-enriched (CD45^+^Ly6G^-^CD19^-^CD20^-^CD3^-/+^TCRgd^-^CD8^-^CD4^+^NK1.1^-^CD25^+^CD103^+^). (K) Absolute number of CD8^+^ T cells (CD45^+^Ly6G^-^CD19^-^CD20^-^CD3^-/+^TCRgd^-^CD8^+^CD4^-^), (L) absolute number of CD8^+^ naïve T cells (CD45^+^Ly6G^-^CD19^-^CD20^-^CD3^-/+^TCRgd^-^CD8^+^CD4^-^CD62L^+^CD44^-^), (M) Absolute number of CD8^+^ Cm T cells (CD45^+^Ly6G^-^CD19^-^CD20^-^CD3^- /+^TCRgd^-^CD8^+^CD4^-^CD62L^+^CD44^+^), (N) absolute number of CD8^+^ Eff T cells (CD45^+^Ly6G^-^CD19^-^CD20^-^CD3^-/+^TCRgd^-^CD8^+^CD4^-^CD62L^-^CD44^+^), (O) absolute number and (P) proportion of of CD8^+^PD-1^+^CX3CR1^+^TIGIT^+^ Eff T cells (CD45^+^Ly6G^-^CD19^-^CD20^-^CD3^-/+^TCRgd^-^ CD8^+^CD4^-^CD62L^-^CD44^+^) and (Q) Dot plot of CD8^+^ Eff T cells for TIGIT, PD1 and CX3CR1 (yellow) of representative mice from each group. Statistics: Mean + SD. Two-way ANOVA with Tukey’s multiple comparison test. * *p* ≤0.05, ** *p* ≤0.01, *** *p* ≤0.001. Cm- central memory; Eff effector T cells.

### 3.6. IL-15 but not IL-15Rα is required to control B16 melanoma expressing NLRC5

The processing and presentation of tumor antigens via the MHC class-I pathway is regulated by the transcriptional activator NLRC5 [51]. The NLRC5 gene is repressed by the polycomb repressor complex [52], which is differentially modulated by the loss of IL-15Rα (Fig. 5E). Therefore, we examined NLRC5 gene expression in MCA tumors and MCA tumor-derived cell lines. The expression of NLRC5 was comparable between WT, *Il15^−/−^* and *Il15ra^−/−^*tumors and cell lines (Fig. 7A). We have shown that B16-F10 tumors expressing NLRC5 (B16-NLRC5) are highly immunogenic and are efficiently controlled in C57Bl/6 mice that is dependent on both CD8^+^ T and NK cells [25, 53]. Therefore, we assessed the growth of B16-NLRC5 tumors in WT, *Il15ra^−/−^* and *Il15^−/−^* mice (Fig. 7B). *Rag1^−/−^Il15^−/−^* mice, used as a control, showed significantly higher tumor growth than WT mice from day 13 post-implantation, that also surpassed tumor growth in *Il15^−/−^* and *Il15ra^−/−^*mice by day 15, pointing towards the contribution of IL-15 signaling-independent tumor control mechanisms of the adaptive immune system. At 18 days post-implantation, tumor growth was significantly higher in *Il15^−/−^*mice than in WT or *Il15ra^−/−^* mice suggesting that IL-15 is required but not trans-presentation by IL-15Rα for the control of tumors with a competent antigen processing and presentation machinery.

### 3.7. Loss of IL-15 or IL-15Rα cause similar paucity of cytolytic lymphoid cells in TILs

Phenotypic analysis of B16-NLRC5 TILs isolated on day 18 post-implantation (using gating strategies shown in Supplementary Figure. 9) showed a significant reduction in CD45^+^ cells in *Il15^−/−^* mice but not in *Il15ra^−/−^* mice compared to WT hosts (Figure. 8A). Among the innate lymphoid cell population, NK, NKT and ILC1-like cells were almost completely absent in the TILs from *Il15^−/−^* and *Il15ra^−/−^*mice, whereas the latter also showed significant reduction in γδ T cells (Fig. 8B-E). B16-NLRC5 TILs from *Il15ra^−/−^* and *Il15^−/−^* mice also contained significantly reduced numbers of CD4^+^ and CD8^+^ T cells when compared to WT mice (Figure. 8F,K). Naïve CD4^+^ and CD8^+^ T cells were minimal in TILs but were higher in DLNs of all three genotypes (Figure. 8G,L, Supplementary Fig. 10). CD4^+^CD25^+^ cells, presumably Tregs, were enriched in TILs compared to DLNs but were comparable in all three genotypes (Fig. 8H, Supplementary Fig. 10). CD4^+^ T central and effector memory cells were reduced in the TILs of *Il15^−/−^* and *Il15ra^−/−^* mice compared to WT, but not statistically significant (Figure. 8I,J). CD8^+^ T central memory (Cm; CD44^+^CD62L^+^) cells and CD8^+^ T effector (Eff; CD44^+^CD62L^-^) cells were significantly reduced in TILs from *Il15ra^−/−^* and *Il15^−/−^* hosts when compared to WT TILs (Figure. 8M,N). However, TILs of *Il15ra^−/−^* mice had significantly more CD8^+^ T EFF cells than *Il15^−/−^* mice. Eff CD8^+^ T of WT and *Il15ra^−/−^* but not *Il15^−/−^*TILs contained a higher proportion of PD-1^+^ CX3CR1^+^TIGIT^+^ cells representing exhausted phenotype but the expression of CX3CR1 suggest that they may not be terminally exhausted, but still maintain effector phenotype (Figure. 8O,P,Q) [52, 54, 55]. Collectively, despite comparable paucity of different cytotolytic innate and adaptive lymphocyte subsets in *Il15ra^−/−^*and *Il15^−/−^* mice, *Il15ra^−/−^* mice controlled B16-NLRC5 tumor growth more efficiently than *Il15^−/−^* hosts (Figure. 7B), indicating that IL-15Rα is dispensable for IL-15-dependent control of tumors with competent antigen processing and presentation machinery.

**Figure 9:**
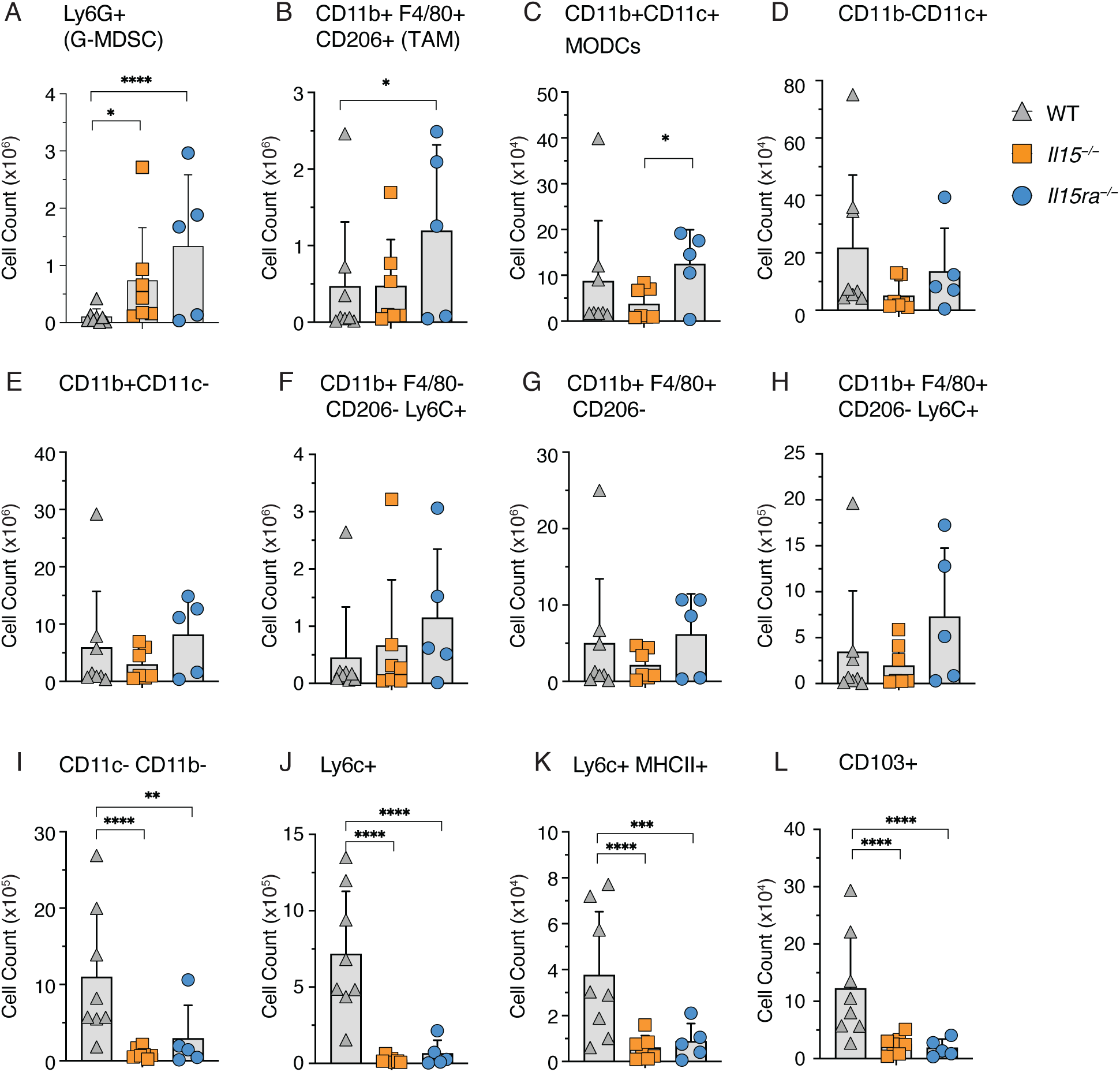
Myeloid subsets in TILs of B16-F10-NLRC5 tumors. Tumor infiltrating lymphocytes (TILs) and single cell suspensions from tumor draining (DLN) and non-tumor draining (NDLN) lymph nodes were isolated at the time of tumor collection from WT, *Il15^−/−^*and *Il15ra^−/−^* mice hosts implanted with B16-F10-NLRC5 cells. Cells were counted and stained with fluorochrome conjugated antibodies for lymphoid and myeloid cell markers and analyzed by flow cytometry using gating strategies depicted in Supplementary Fig. S9. Data from 7-9 mice per group from three independent experiments are shown. The cell numbers in TILs were equalized to the weight of the tumor mass (TILs per gram; TILS(g^-1^)). (A) Absolute number of Ly6G^+^G-MDSC (CD45^+^Ly6G^+^Ly6C^lo^CD11b^+^), (B) Absolute number of TAMs (CD45^+^Ly6G^-^CD19^-^CD20^-^CD3^+/-^TCRgd^-^CD4^-^CD8^-^NK1.1^-^CD11b^+^CD11c^-^ F4/80^+^CD206^+^), (C) Absolute number of MODCs (CD45^+^Ly6G^-^CD19^-^CD20^-^CD3^+/-^TCRgd^-^CD4^-^CD8^-^NK1.1^-^CD11b^+^CD11c^+^), (D) Absolute number of CD11c^+^DCs (CD45^+^Ly6G^-^CD19^-^CD20^-^ CD3^+/-^TCRgd^-^CD4^-^CD8^-^NK1.1^-^CD11b^-^CD11c^+^), (E-H) Absolute number of monocytes (CD45^+^Ly6G^-^CD19^-^CD20^-^CD3^+/-^TCRgd^-^CD4^-^CD8^-^NK1.1^-^CD11b^+^CD11c^-^) expressing or not F4/80, CD206 and Ly6C and (I-L) Absolute number of CD11c^-^CD11b^-^ immature monocytes (CD45^+^Ly6G^-^CD19^-^CD20^-^CD3^+/-^TCRgd^-^CD4^-^CD8^-^NK1.1^-^CD11b^-^CD11c^-^) expressing or not Ly6C, MHCII and CD103. Statistics: Mean + SD. Two-way ANOVA with Tukey’s multiple comparison test. * *p* ≤0.05, ** *p* ≤0.01, *** *p* ≤0.001.

### 3.8. Lack of IL-15 or IL-15Rα differentially impact myeloid cell subsets of TILs

Characterization of myeloid lineage cells in B16-NLRC5 TILs revealed a significant enrichment of granulocytic-myeloid-derived suppressor cells (G-MDSC: Ly6G^+^Ly6C^lo^CD11b^+^) in *Il15^−/−^*and *Il15ra^−/−^* mice compared to WT controls, and elevated numbers of tumor associated macrophages (TAMs: CD11b^+^F4/80^+^CD206^+^) in TILs of *Il15ra^−/−^* mice (Fig. 9A,B). On the other hand, monocyte-derived MoDCs (CD11b^+^CD11c^+^) and CD11c^+^ DCs in TILs of *Il15^−/−^* and *Il15ra^−/−^* mice were comparable to WT mice (Fig. 9C,D). The number of monocytes (CD11b^+^) was comparable in TILs of WT, *Il15^−/−^* and *Il15ra^−/−^*mice (Fig. 9E-H). However, CD11c^−^CD11b^−^ immature monocytes that express Ly6C, MHC-II or CD103 were significantly reduced in TILs of both *Il15^−/−^* and *Il15ra^−/−^* mice (Fig. 9I-L). These immature monocyte precursors, which are mobilized during inflammatory conditions, are considered pro-resolving macrophages [56]. These data suggest that IL-15 signaling restrains infiltration of G-MDSCs, TAMs and immature monocyte precursors into immunogenic tumors, independently of trans-presentation by IL-15Rα, which may also contribute to IL-15-driven anti-tumor immune responses.

## 4. Discussion

Since the cloning of IL-15 a little more than 30 years ago, our collective understanding of the structure, functions and therapeutic applications of IL-15 has resulted in the approval of IL-15-derived biologics in cancer immunotherapy [57]. IL-15 is used in combination with other immunotherapeutic approaches to activate NK and CD8^+^T cells to control tumor growth. While our knowledge on the IL-15 mediated activation of NK and CD8^+^T cells is extensive, other aspects of IL-15 biology is not well understood. Most of the studies related to IL-15 and the development of IL-15 mimics for tumor immunotherapy have centered around the ability of IL-15 to activate ILC1, NK, γδ T and NKT and CD8^+^ T cells, all of which requires trans-presented IL-15 [15]. Tissue-specific deletion of IL-15Rα revealed the requirement for localized trans-presentation by IL-15Rα for the survival of tissue-resident IL-15-dependent lymphocyte subsets [58, 59]. Thus, the availability of circulating IL-15 does not replace the membrane bound IL-15:IL-15Rα in the maintenance of lymphoid populations in tissues. Additionally, IL-15 administration induces cytokine storm despite its short half-life [60]. On the other hand, soluble IL-15Rα that is cleaved from the cell surface acts as an antagonist of IL-15 signaling as it competes with membrane bound IL-15Rα for IL-15 [16]. Other than trans-presenting IL-15 to IL-15Rβ:γ_c,_ IL-15Rα has no identified functions as it does not contain any signaling modules. Taking into consideration the various constraints associated with systemic toxicity and the complexity of IL-15 signaling, approaches to the development of anti-cancer immunotherapeutic agents have evolved over the past 3 decades and the first biologic – N-803 – has been approved for localized intravesicular administration as part of combination therapy with BCG for non-muscle invasive bladder cancer (NMIBC) in 2024 [57, 61].

Various lines of evidence from the literature indicate that IL-15 signaling independent of IL-15Rα mediated trans-presentation plays important roles in controlling infections and autoimmunity [5, 8, 14, 16]. IL-15 activates the innate immune system during infections and hence, is indispensable for controlling infections [15], but we showed that trans-presented IL-15 was not essential to control Listeria and Salmonella infections [14]. In autoimmunity, IL-15 deficient mice are protected from autoimmune type 1 diabetes and certain forms of psoriasis [8, 16], but not IL-15Rα deficient mice [5]. In psoriasis, absence of IL-15Rα increased the disease severity while administration of soluble IL-15Rα reduced inflammation and improved the clinical status [16]. In contrast in models of Th-17 mediated autoimmune diseases like EAE, absence of IL-15 exacerbated the severity, as IL-15 can suppress Th17 development under certain circumstances [62, 63]. Thus, in the above models, the mechanisms by which IL-15 and/or trans-presented IL-15 contributes to each of the above pathologies is distinct. The role of IL-15Rα mediated trans-presentation in mediating IL-15-driven antitumor immune responses remains poorly understood. Comparable growth of three different established tumor cell lines in WT, IL-15 deficient and IL-15Rα deficient mice indicate that the absence of IL-15-signaling directly or through trans-presentation for the maintenance of IL-15 dependent immune subsets do not impact tumor growth, probably due to the aggressivity of the tumor cell lines. In these models, innate lymphocyte subsets play a minimal role in controlling growth of solid tumors [34]. Constant surveillance by cytotoxic innate immune cells plays an important role in hematological malignancies and viral or oncogene-associated cancers, but not in MCA-tumor model [46, 64]. Studies using depleting anti-NK1.1 and anti-NKG2D antibody to determine the relative contribution of innate lymphocyte subsets, such as NK, NKT, γδ T cells and ILCs in immune surveillance of solid tumors is not very clear as treatment will also deplete specific subsets of effector CD8^+^ T cells that express these markers [65–71]. Deficiency in Rag1, perforin, IFNγ, IFNγR or trail that severely diminishes the cytotoxicity of CD8^+^ T cells compromises immune surveillance, resulting in early onset of MCA-induced tumors [42, 72–75]. Our data show that absence of IL-15 shows negligible impact on the incidence of MCA-induced tumors, indicating that IL-15 is dispensable for immunosurveillance of endogenously arising solid tumors. Nonetheless, the delayed tumor growth or the lack of tumor formation by MCA tumor-derived cell lines from *Il15^−/−^* and *Il15ra^−/−^* mice in all the WT recipients suggest that both IL-15 and IL-15Rα contribute to tumor immunoediting, possibly from their shared positive impact on CD8+ T cell-mediated immune responses.

Strikingly, the reduced incidence of spontaneous MCA-induced tumors in *Il15ra^−/−^* mice when compared to *Il15^−/−^* or WT mice reiterates two possible non-mutually exclusive mechanisms: (a) IL-15Rα restrains antitumor immune responses that may include both IL-15-driven and non-IL-15-mediated responses, and (b) enhancement of IL-15Rα-independent IL-15 signaling in the absence of IL-15Rα-mediated trans-presentation. Our efforts to understand the mechanisms using proteomic approaches shed some light on the potential mechanisms involved. Notably, MCA-induced tumors from IL-15Rα deficient mice showed lower levels of proteins implicated in neutrophil net formation (NET), which can be a reflection of reduced NET-induced immune suppression, and hence better tumor responses [76]. In addition, pathways associated with chemical carcinogenesis -ROS, was reduced in MCA-tumor proteome from *Il15ra^−/−^* mice when compared to WT and *Il15^−/−^*mice. The high metabolic activity of tumor cells leads to increased ROS production that contributes to cancer progression through many pathways [77]. However, it will be simplistic to assume that these pathways that are differentially regulated are intrinsic to the tumors. Comparison of proteomes of MCA tumors and MCA tumor derived cell lines shows that many of the changes in the immune related pathways can be attributed to the host immune system rather than the to the tumors themselves. Thus, IL-15Rα seems to impact on tumor associated signaling pathways that are both tumor intrinsic and modulated by the tumor microenvironment.

Poor immunogenicity of tumors coupled with immunosuppressive tumor environment is a basic feature of tumors with poor prognosis. One of the approaches involves enhancing tumor antigen presentation by expressing the MHC-I trans-activator NLRC5 in tumors [30, 53, 78, 79]. NLRC5-expressing B16-F10 tumors were well controlled not only in WT mice as we have shown before [25], but also in IL-15Rα deficient mice. These results are surprising as TILs from both IL-15 deficient and IL-15Rα deficient mice had reduced numbers of NK, NKT, ILC1-like and CD8^+^ T cells, including the different CD8^+^ T cell subsets. These observations open up the possibility that tumor intrinsic mechanisms play a greater role in efficient anti-tumor immunity. This is further supported by our previous observations that mice that controlled NLRC5-expressing B16 tumors, were able to reject challenge with parental B16 cell line, possibly due to the expanded repertoire of tumor derived peptides that are presented [25, 53, 78]. Thus, therapies that target tumors to become immunogenic may be more effective than trying to target the immune system directly. This point of view may be intuitive as the tumors grow as close to being ‘self’ as much as possible that the immune system ignores the cold tumors. Breaking this tolerance needs the tumor to become foreign through a plethora of new antigens, and preferably accompanied by innate immune stimulation. Thus, mechanisms that expand the tumor-associated antigen repertoire, rather than targeting activation of specific subsets of immune cells alone, hold promise for developing more efficient antitumor immune therapies.

## Acknowledgements

We thank the proteomics platform and the flowcytometry platform, Université de Sherbrooke for data acquisition.

## Funding

This work was supported by CIHR grants to SR and SI (PJT- 173482 and PJT-203990). HA-C is Junior 2 Clinical Research Scholar from the Fonds de recherche du Québec-Santé and holds the André-Lussier research chair of the Université de Sherbrooke. MM and SK are recipients of PhD scholarships from Université de Sherbrooke.

## Authors’ contributions

Conceptualization, S.R., S.I., F.R.; Methodology, F.R., S.A.A., A.S., M.M., S.K., E.D., A.A.C. and S.R.; formal analysis, F.R., S.A.A., M.M., S.K., J.-F.L., H.A.C, S.I. and S.R.; resources, F.-M.B., S.R. and S.I.; data curation, F.R., M.M., J.-F.L. and S.R.; writing—original draft preparation, F.R. and S.R.; writing—review and editing, F.R., M.M., J.-F.L., S.I. and S.R.; supervision, S.R. and S.I.; funding acquisition, S.R and S.I.

## Conflict of interest

The authors declare that they have no competing financial interests.

## Notes

### Competing Interest Statement

The authors have declared no competing interest.

