## Supplementary table 1 for "Trans-presentation of IL-15 by IL15Rα attenuates tumor immune surveillance and is dispensable for IL-15-dependent tumor growth control"

Rexhepi F., et al.,

### Supplementary Tables

#### Supplementary Table S1. List of primers used for RT-qPCR.

| Gene name | Gene ID | Sense primer | Anti-sense primer | Amplicon Size (bp) |
|---|---|---|---|---|
| *Cxcl9* | NM_008599.4 | AGTCCGCTGTTCTTTTCCTC | TGAGGTCTTTGAGGGATTTGTAG | 141 |
| *Cxcl10* | NM_021274.2 | CCAAGTGCTGCCGTCATTTTC | GGCTCGCAGGGATGATTTCAA | 157 |
| *Cd8a* | NM_001081110.2 | CATCACTCTCATCTGCTACCAC | TTTTCTCTGAAGGTCTGGGC | 98 |
| *Cd4* | NM_013488.3 | GTTCGGCATGACACTCTCAG | CCTTCTCTGCCTTCCACATC | 146 |
| *Klrb1c* (NK1.1) | NM_001159904.2 | TGGCACAACTTTCAATTCTGATG | GGTTTAGTTCCTTTTGGCAGATC | 140 |
| *Cd274* (PD-L1) | NM_021893.3 | CTCATTGTAGTGTCCACGGTC | ACGATCAGAGGGTTCAACAC | 149 |
| *Ifng* | NM_008337.4 | CCTAGCTCTGAGACAATGAACG | TTCCACATCTATGCCACTTGAG | 150 |
| *Ifnb1* | NM_010510.2 | TGTCCTCAACTGCTCTCCAC | CTGAAGATCTCTGCTCGGACC | 112 |
| *Klrd1* | NM_010654.5 | TCAACACCTTCTCCAACCAC | CTGATGCCCAACCCACTT | 84 |
| *Klrc2* | NM_010653.5 | AATCTCTTTCACTGGTCTCATGG | GCAAATTCATCTAAAGGGAGCC | 262 |
| *Klrk1* | NM_033078.4 | GCTGGTTAAGTCCTATCACTGG | TTGAGCCATAGACAGCACAG | 143 |
| *36B4 (Rplp0)* | NM_007475.5 | TCTGGAGGGTGTCCGCAAC | CTTGACCTTTTCAGTAAGTGG | 154 |

#### Supplementary Table S2. List of antibodies used on a Cytoflex 30 flow cytometer.

| **Antibody** | **Fluorophore** | **Clone** | **Company** | **Cat#** |
|---|---|---|---|---|

| | | | | |
|---|---|---|---|---|
| Flexible Viability Dye | eFluor 780 | - | eBiosciences | 65-0865-18 |
| CD45 | Brilliant Violet 605 | 30-F11 | Biolegend | 103140 |
| CD19 | PE | 6D5 | Biolegend | 115508 |
| NK1.1 | APC-Cy7 | PK136 | Biolegend | 108724 |
| TCRb | PE/Dazzle594 | H57-597 | Biolegend | 109240 |
| CD8a | eFluor450 | 53-6.7 | eBiosciences | 48-0081-82 |
| CD4 | AlexaFluor700 | GK1.5 | eBiosciences | 50-168-51 |
| CD62L | APC | MEL-14 | eBiosciences | 17-0621-83 |
| CD44 | FITC | IM7 | Biolegend | 103006 |
| CD69 | PECy7 | H1.2F3 | Biolegend | 104512 |
| CD279 (PD-1) | BV785 | 29F.1A12 | Biolegend | 135225 |
| TCRgd | PerCP | GL3 | Biolegend | 118117 |
| CD45 | Brilliant Violet 510 | 30-F11 | Biolegend | 742863 |
| Ly6G | PerCP | 1A8 | Biolegend | 127654 |
| CD11c | Alexa Fluor 700 | N418 | eBiosciences | 56-0114-82 |
| CD11b | eFluor450 | M1/70 | eBiosciences | 48-0112-82 |
| MHC II | PECy7 | M5/114.15.2 | Biolegend | 107630 |
| Ly6C | Alexa Fluor 647 | HK1.4 | Biolegend | 128016 |
| F4/80 | Brilliant Violet 605 | BM8 | Biolegend | 123133 |
| CD122 | FITC | TM-B1 | eBiosciences | 11-1222-82 |
| CCR2 | PE | SA203G11 | Biolegend | 150610 |
| CX3CR1 | PE/Dazzle 594 | SA011F11 | Biolegend | 149013 |

**Supplementary Table S3. List of antibodies used on a BD FACSDiscover™ S8 cell sorter.**

| **Antibody** | **Fluorophore** | **Clone** | **Company** | **Cat#** |
|---|---|---|---|---|
| CD197 (CCR7) | BV605 | 4B12 | BD | 740431 |
| CX3CR1 | BB700 | Z8-50 (Z8-50.23) | BD | 567812 |
| CD185 (CXCR5) | RB613 | 2G8 | BD | 759651 |
| CD194 (CCR4) | PE-Cy7 | 2G12 | BioLegend | 131214 |
| CD183 (CXCR3) | PE-Fire640 | S18001A | BioLegend | 155922(BLG) |
| CD197 (CCR7) | BV605 | 4B12 | BD | 740431 |
| CD69 | BUV805 | 30-F11 | BD | 741927 |
| Ly6G | BUV737 | 1A8 | BD | 568346 |
| Ly49A | BUV661 | A1 | BD | 749810 |
| CD27 | BUV615 | LG.3A10 | BD | 751529 |
| NK1.1 | BUV563 | PK136 | BD | 741233 |
| CD45 | BUV496 | 145-2C11 | BD | 569673 |
| Live/Dead (FVS440 UV) | FVS440UV | - | BD | 566332 |
| XCR1 | Spark UV 387 | ZET | BioLegend | 148236(BLG) |
| CD62L | BUV395 | MEL-14 | BD | 569400 |
| MHC-I | BV786 | BD | AF6-88.5 | 742863 |
| CD279 (PD-1) | BV750 | 29F.1A12 | BD | 568635 |
| CD107a | BV711 | 1D4B | BD | 564348 |
| CD64 A/B | BV650 | X54-5/7.1 | BD | 740622 |
| CD11c | BV510 | HL3 | BD | 562949 |
| CD44 | BV480 | IM7 | BD | 566116 |

| | | | | |
|---|---|---|---|---|
| CD4 | RV828 | GK1.5 | | 5726748 |
| TIGIT | BV421 | 1G9 | BD | 565270 |
| CD152 (CTLA-4) | PE | BNI3 | BD | 553720 |
| CD11b | PerCP-Fire806 | M1/70 | BioLegend | 101294 |
| Singlec-H | RB780 | 440c | BD | 755362 |
| CD103 | RB744 | M290 | BD | 758108 |
| TCRgd | RB705 | GL3 | BD | 757547 |
| CRIg (VSIG4) | RB670 | 17C9 | BD | 771636 |
| CD49a | RB545 | Ha31/8 | BD | 756784 |
| CD25 | BB515 | PC61 | BD | 564424 |
| I-A/I-E (HLA-DR) | PE-Fire810 | M5/114.15.2 | BioLegend | 107667 |
| CD206 | PE-Fire700 | C068C2 | BioLegend | 141742(BLG) |
| IgD | PE-Cy5 | 11-26c-2a | BD | 624350 |
| TIM-4 | RY610 | 21H12 | BD | 758786 |
| CD8a | RY743 | 53-6.7 | BD | 572203 |
| CD19 | RY586 | 1D3 | BD | 753680 |
| CD20 | APC-Fire810 | SA275A11 | BioLegend | 150438 |
| Ly-6C | APC-Cy7 | AL-21 | BD | 560596 |
| CD274 (PD-L1) | R718 | 10F.9G2 | BD | 568585 |
| CD3e | APC | H1.2F3 | BD | 565643 |
| F4/80 | AF647 | T45-2342 | BD | 565853 |

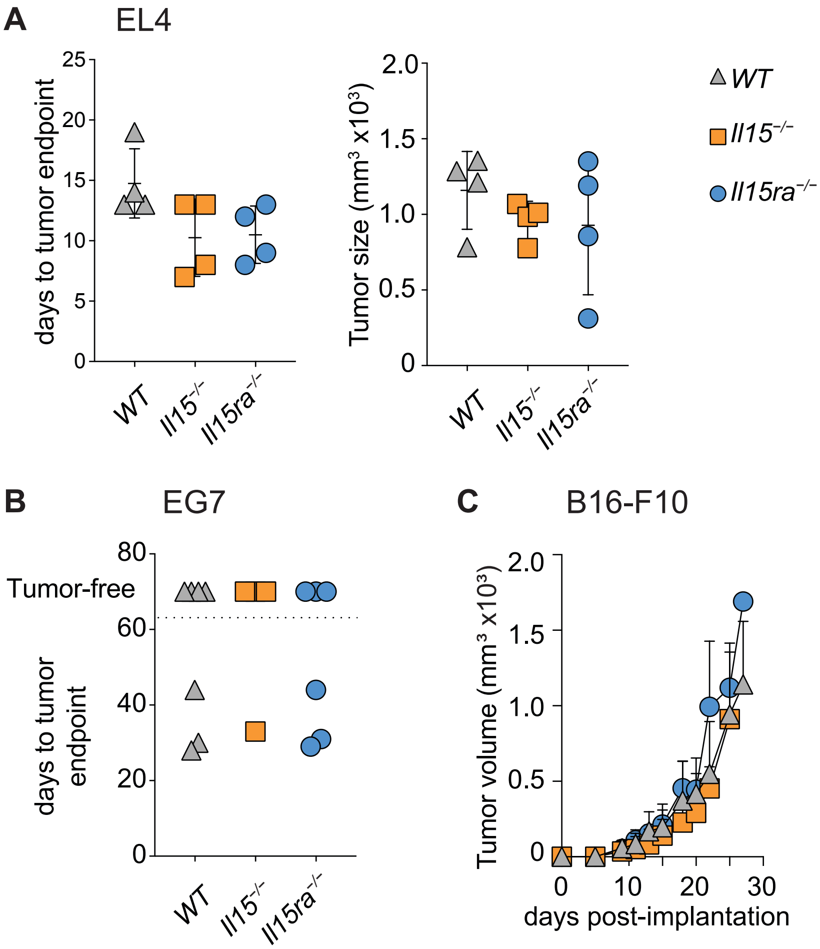


**Supplementary Figure S1. Loss of IL-15 or IL-15R$\alpha$ does not affect the growth of established tumor cells lines.** WT, *Il15*$^{-/-}$ and *Il15ra*$^{-/-}$ mice were implanted in their right flank with (A) EL4 lymphoma or (B) EG7 cells (EL4 expressing chicken ovalbumin) or (C) B16-F10 melanoma and tumor growth was measured every 2 days from day 8 onwards. When the tumor reached the endpoint (20 mm in diameter in any one direction) in more than one of the recipient mice, all mice were euthanized. 4-7 mice per group.

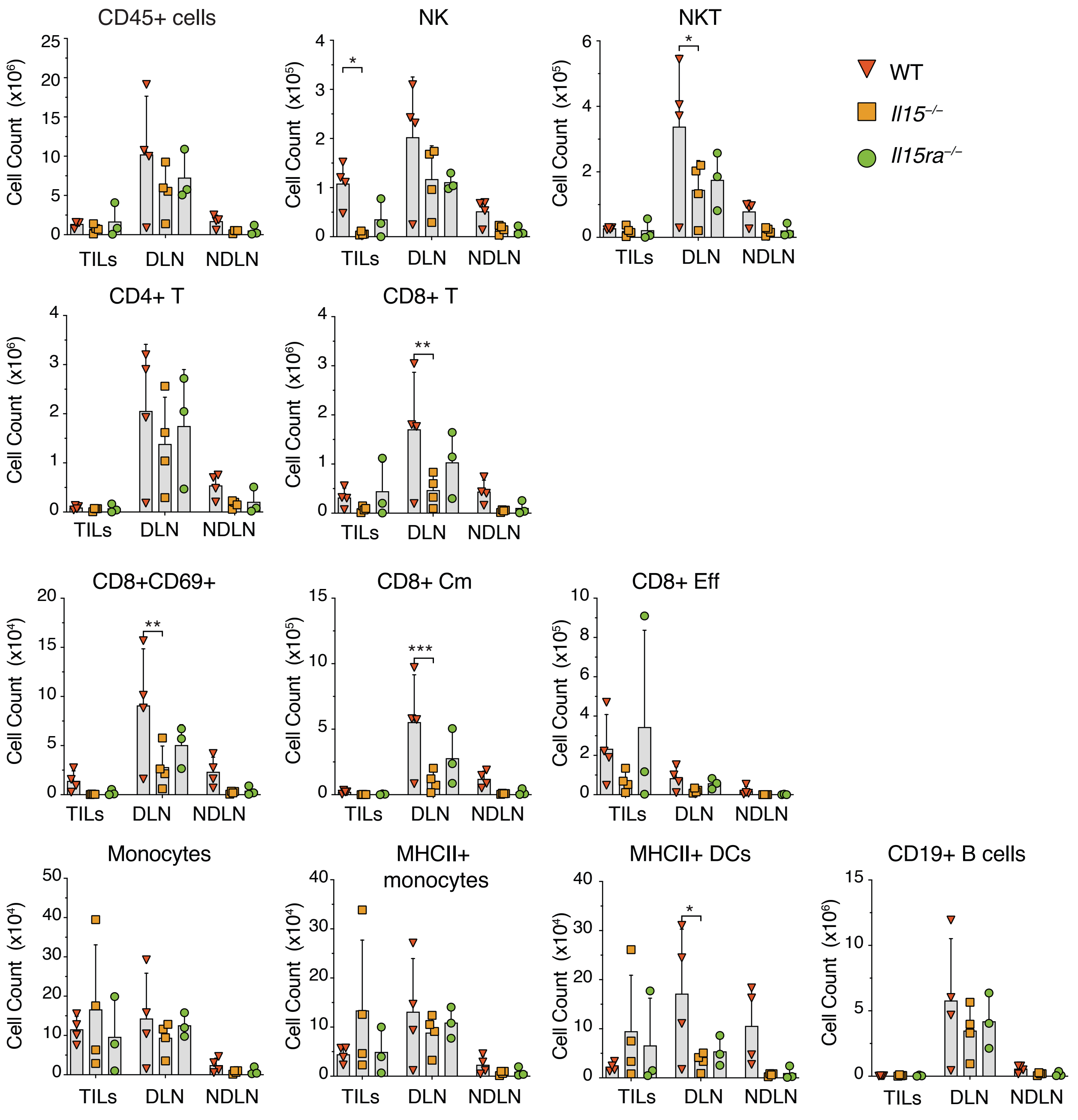


**Supplementary Figure S2. Immune cell infiltration in B16 tumors from WT, *Il15*$^{-/-}$ and *Il15ra*$^{-/-}$ mice.** Tumor infiltrating lymphocytes (TILs) and single cell suspensions from tumor draining (DLN) and non-tumor draining (NDLN) lymph nodes were isolated at the time of tumor collection from WT, *Il15*$^{-/-}$ and *Il15ra*$^{-/-}$ mice hosts implanted with B16-F10 melanoma cells. Cells were counted and stained with fluorochrome-conjugated antibodies for lymphoid and myeloid cell markers and analyzed by flow cytometry. Data from 4 mice per group from two independent experiments are shown. The cell numbers in TILs were equalized to the weight of the tumor mass (TILs per gram; TILS($g^{-1}$)). Absolute number of the indoicated immune cell populations [Leukocytes ($CD45^+$), NK cells ($NK1.1^+TCR\beta^-$), NKT cells

(NK1.1$^+$TCRβ $^+$) cells, CD4$^+$ T cells CD8$^+$ T cells, CD8+ T cell subsets (CD8$^+$ Cm– CD8$^+$CD44$^+$CD62L$^+$ and CD8$^+$ Eff – CD8$^+$CD44$^+$CD62L$^-$), monocytes (CD11b$^+$Ly6C$^+$), DCs (CD11b$^-$CD11c$^+$) and B cells (CD19$^+$)] were calculated from their percentages within the gated populations. Statistics: Mean + SD. Two-way ANOVA with Tukey's multiple comparison test. * $p \leq 0.05$, ** $p \leq 0.01$, *** $p \leq 0.001$. Abbreviations: CM, central memory; EFF effector; DC, dendritic cells.

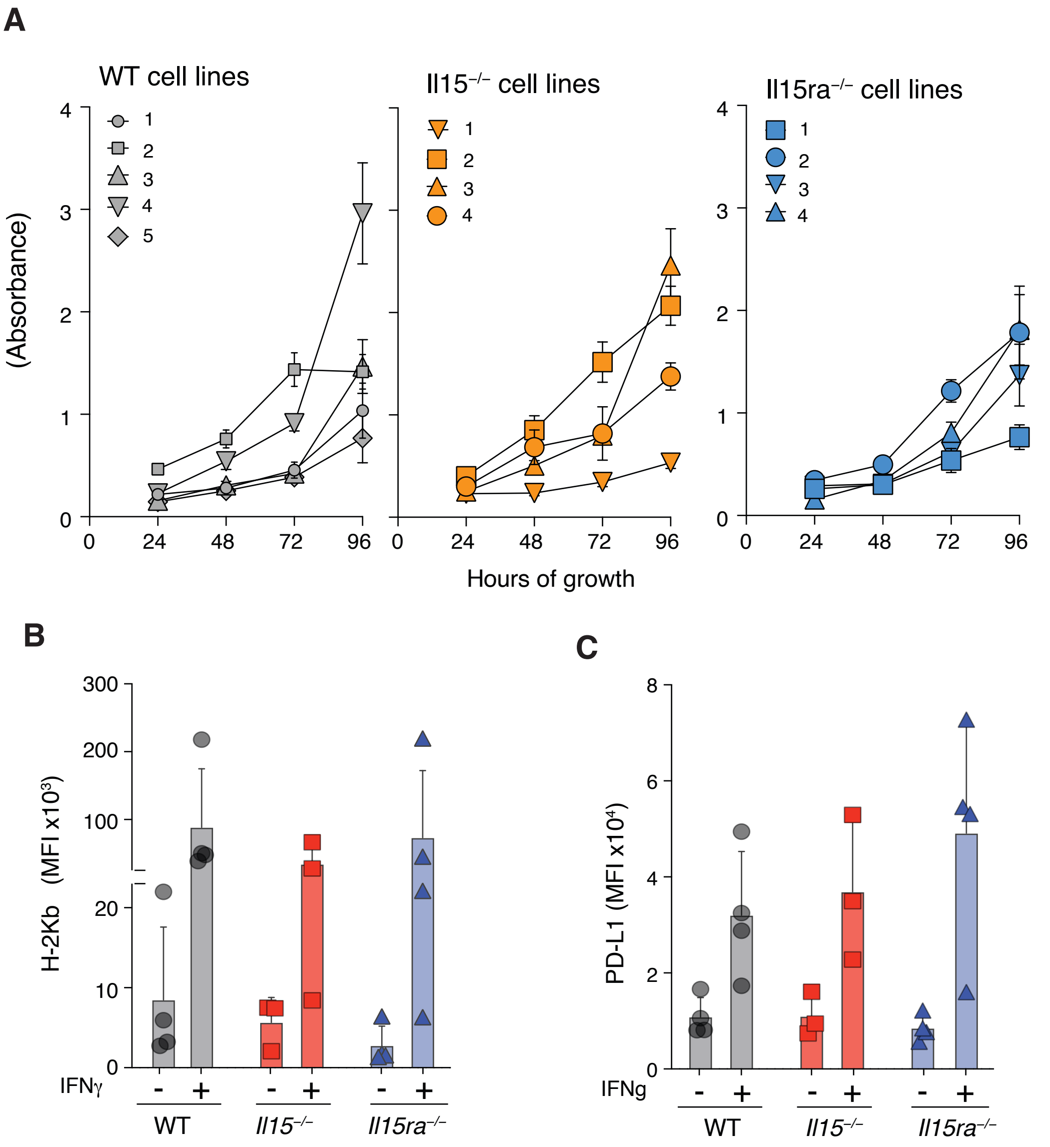


**Supplementary Figure S3. Comparable growth of MCA tumor-derived cell lines *in vitro*.** (A) 4-5 independent cell lines were established from the tumors *in vitro*. Growth of the different MCA-tumor derived cell lines *in vitro* as determined by MTT assay. (B,C) Cell lines were exposed to mouse IFNγ (4ng/ml) for 18 h) and the induction of MHC-I (*H2kb*) and PD-L1 (*Cd274*) was evaluated by RT-qPCR.

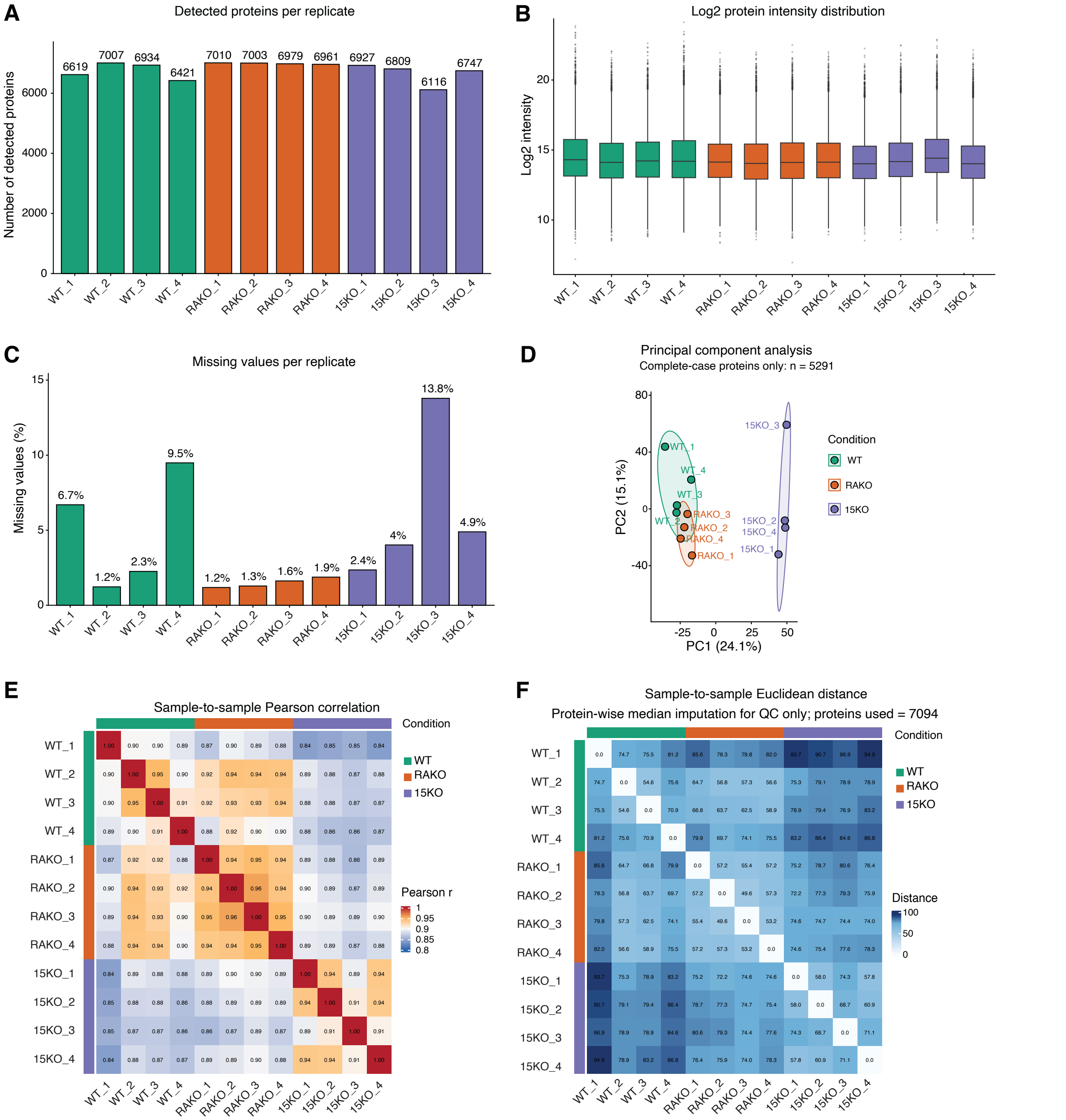


**Supplementary Figure S4. Quality control of mass spectrometry data on MCA tumor proteomes**. Quadruplicate samples of proteins extracted from WT, *Il15*$^{-/-}$ (15KO) and *Il15ra*$^{-/-}$ (RAKO) tumors were subjected to shotgun proteomics using LC-MS/MS. (A) Number of individual proteins identified in each tumor by more than one peptide; (B) protein intensity distribution in the samples; (C) Missing value patterns across samples; (D) Principal component analysis (PCA) plot; (E) Pearson correlation between samples; (F) Sample to sample Euclidean distance.

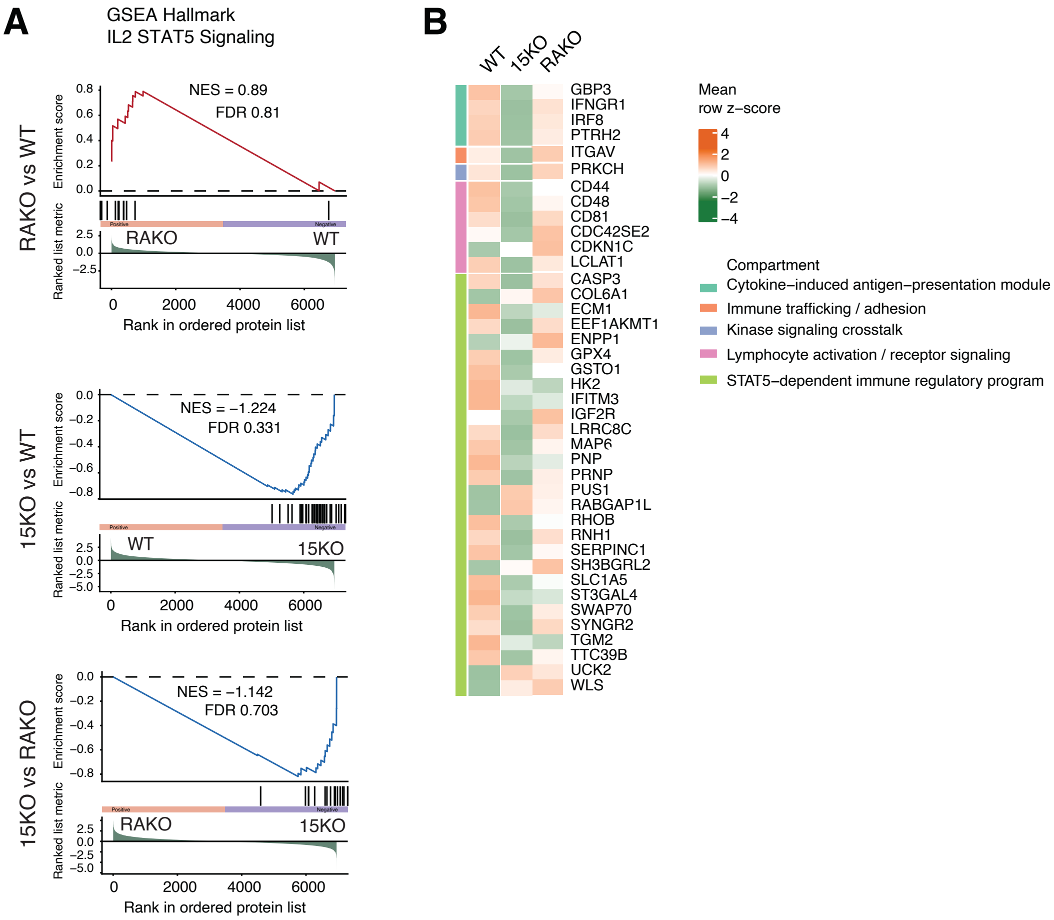


**Supplementary Figure S5. Differential enrichment of IL-2-Stat5 signaling pathway proteins in WT, *Il15ra*$^{-/-}$ and *Il15*$^{-/-}$ tumors.** (A) Unbiased GSEA of Hallmark IL-2 Stat5 signaling pathway proteins that are differentially expressed in WT, *Il15ra*$^{-/-}$ and *Il15*$^{-/-}$ tumors. (B) Heatmap representation of normalized abundance levels of the indicated proteins (row Z-score, color coded) within the functional components of IL-2 Stat5 signaling pathway (color coded and indicated) in quadruplicate samples of WT, *Il15ra*$^{-/-}$ and *Il15*$^{-/-}$ tumors.

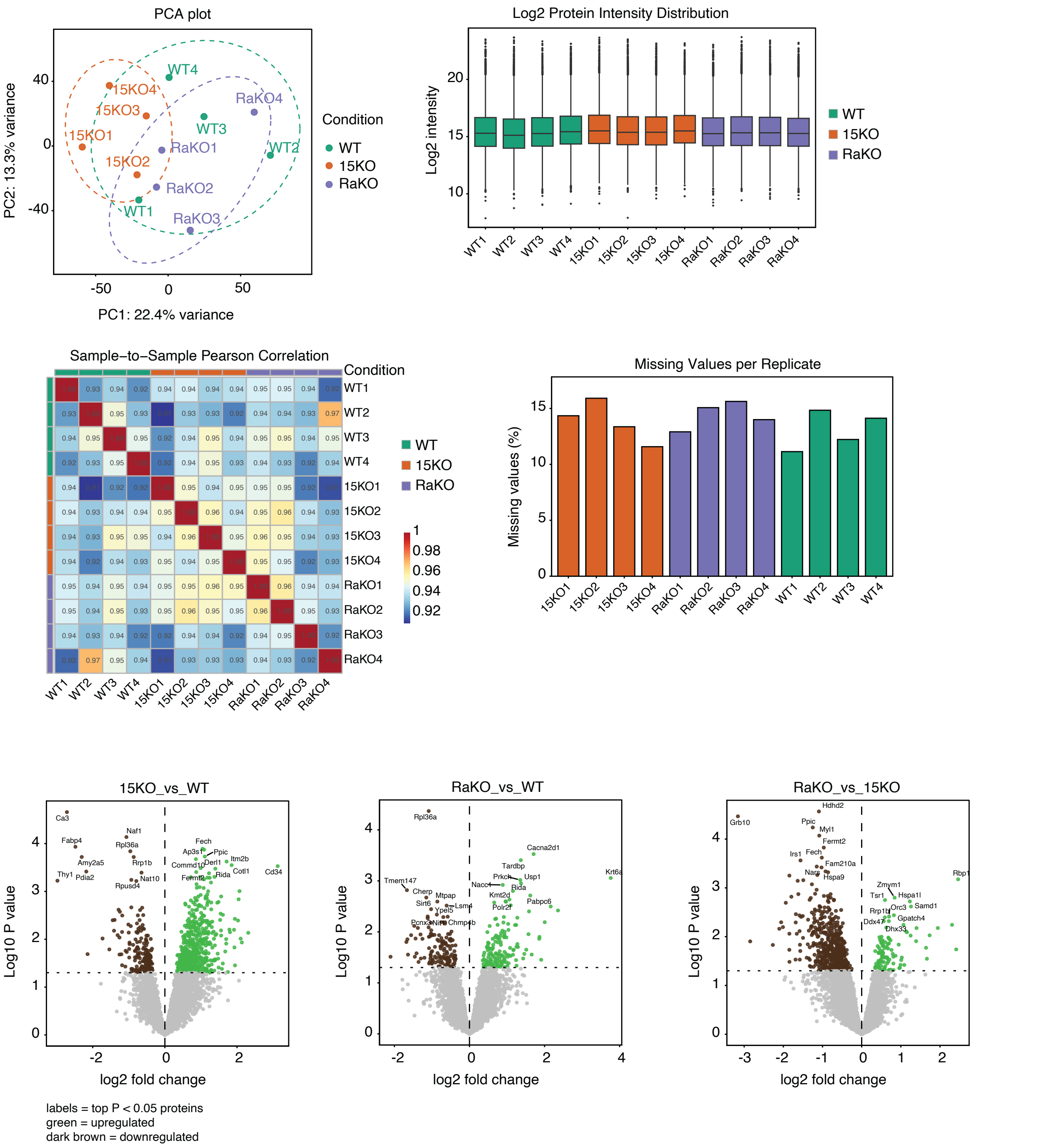


**Supplementary Figure S6. Quality control of mass spectrometry data on proteomes of MCA tumor derived cell lines**. Quadruplicate samples of proteins extracted from WT, *Il15ra*$^{-/-}$ and *Il15*$^{-/-}$ MCA tumor-derived cell lines were subjected to LC-MS/MS. Principal component analysis (PCA) plot, protein intensity distribution in the samples, Pearson correlation between samples, missing value patterns across samples and volcano plot of proteins detected in IL-

15KO and IL-15R$\alpha$KO compared to WT cell lines, and IL-15KO compared to IL-15R$\alpha$KO cell lines are shown.

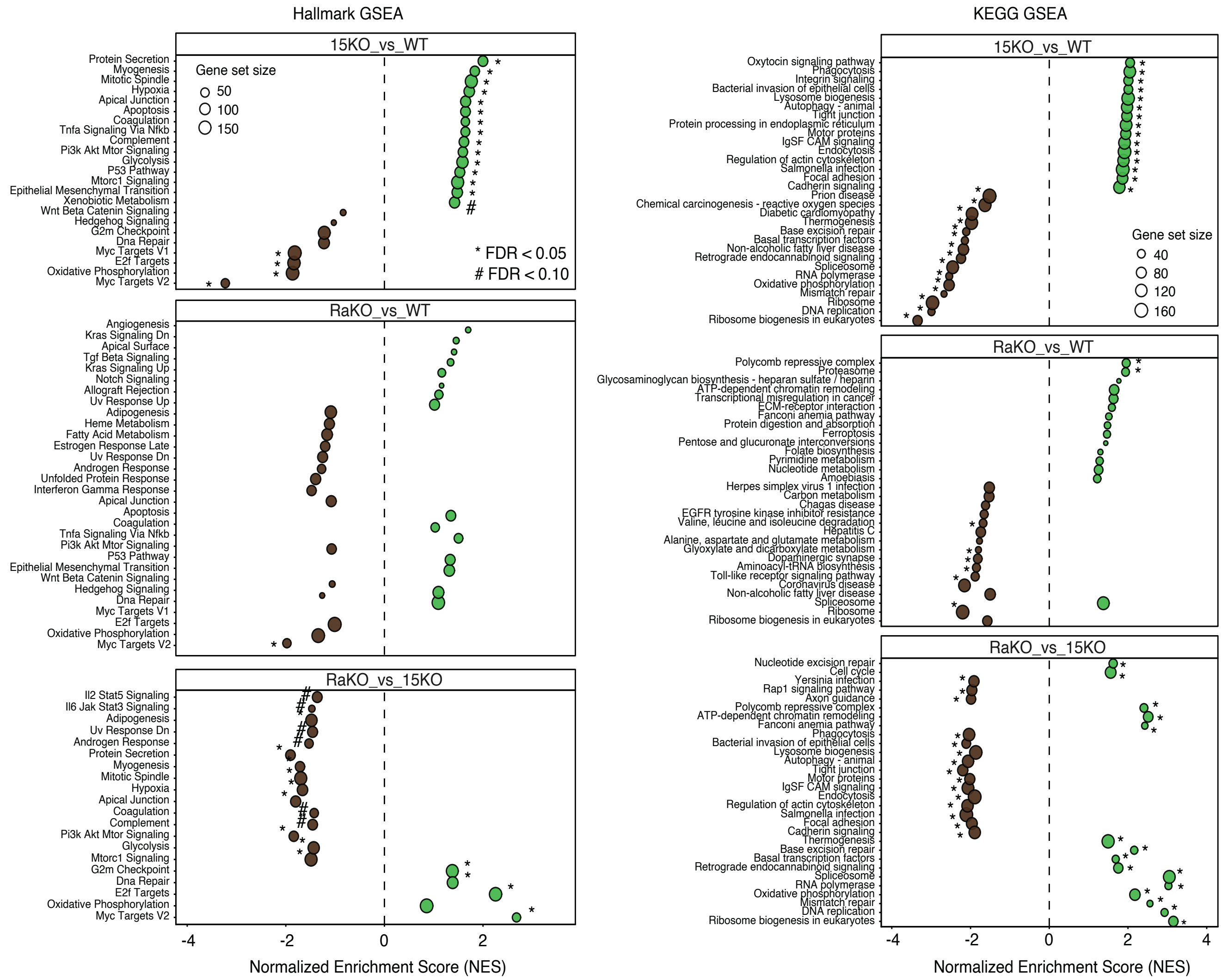


**Supplementary Figure S7: Differentially expressed pathways from proteome analyses of MCA tumor-derived cell lines.** Pathways that are different between IL-15KO and IL-15R$\alpha$KO compared to WT cell lines, and IL-15KO compared to IL-15R$\alpha$KO cell lines by Hallmark and KEGG GSEA analyses are shown.

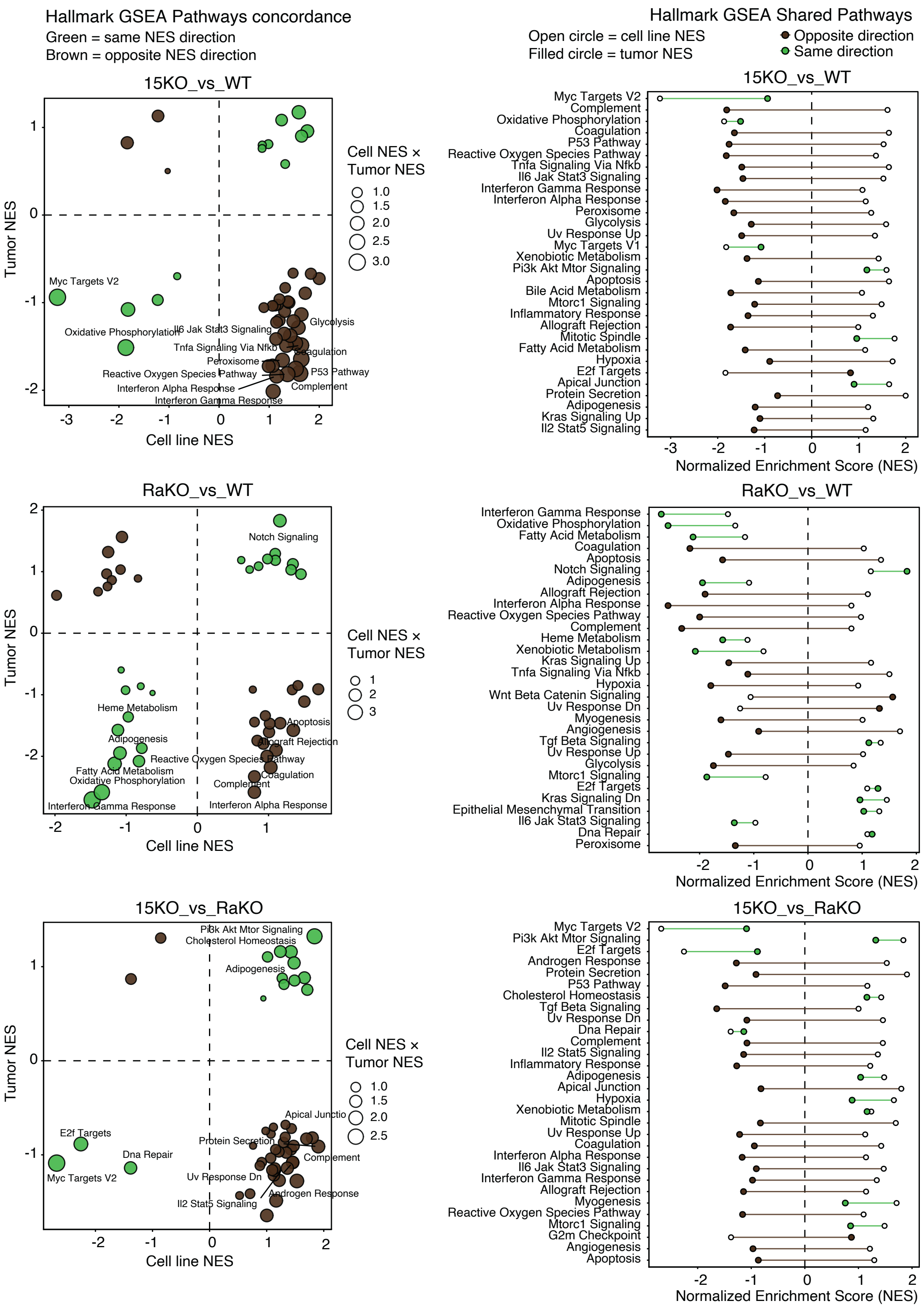


**Supplementary Figure 8: Comparison of differentially expressed pathways from proteome analyses of MCA tumors and MCA tumor derived cell lines.** Hallmark Pathway concordance and shared pathways are shown.

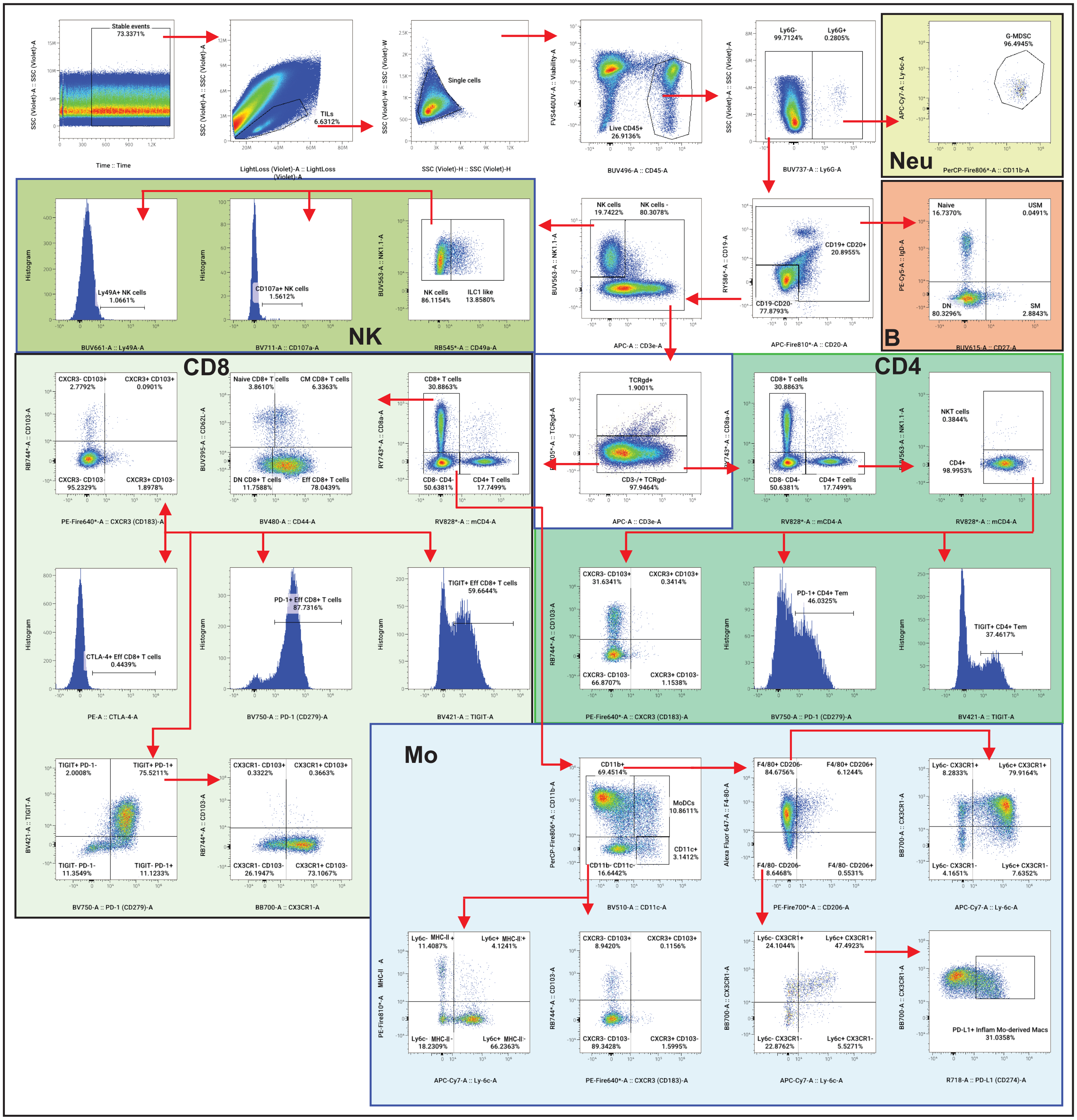

**Supplementary Figure S9: Gating strategy for data shown in Fig. 8 and 9.** Arrows indicate the gating sequence. Single cells were gated out for dead cells followed by selection of CD45+ cells. Within CD45+ cells, neutrophils, B cells, NK cells and gd T cells were gated out sequentially. CD4+ and CD8+ T cells were gated out in the remaining population. Monocytes were selected on the CD4-CD8- cells.  Abbreviations: Mo, monocyte/macrophage subsets. Neu, neutrophils.

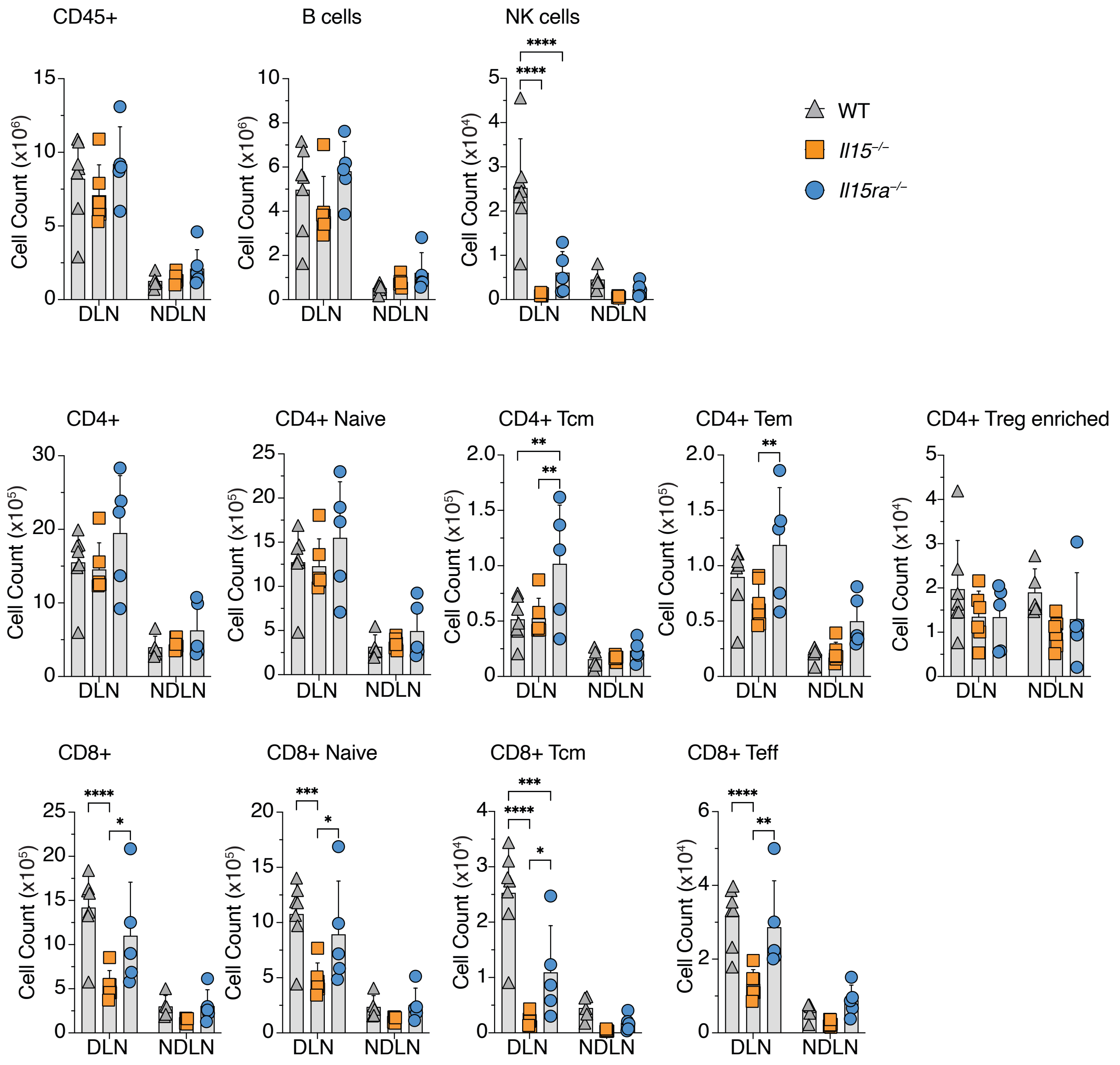


**Supplementary Figure S10. Distribution of select immune subsets detected in the draining and non-draining lymph nodes of B16-NLRC5 tumor bearing mice** (data complementary to Figures 8 and 9) Statistics: Mean + SD. Two-way ANOVA with Tukey's multiple comparison test. * $p$ ≤0.05, ** $p$ ≤0.01, *** $p$ ≤0.001. Cm, central memory; Eff, effector T cells.
